# Miro1-dependent Mitochondrial Dynamics in Parvalbumin Interneurons

**DOI:** 10.1101/2020.10.06.328534

**Authors:** Georgina Kontou, Pantelis Antonoudiou, Marina Podpolny, I. Lorena Arancibia-Carcamo, Nathalie F. Higgs, Blanka R. Szulc, Guillermo Lopez-Domenech, Patricia C. Salinas, Edward O. Mann, Josef T. Kittler

## Abstract

The spatiotemporal distribution of mitochondria is crucial for precise ATP provision and calcium buffering required to support neuronal signaling. Fast-spiking GABAergic interneurons expressing parvalbumin (PV) have a high mitochondrial content reflecting their large energy utilization. The importance for correct trafficking and precise mitochondrial positioning remains poorly elucidated in inhibitory neurons. Miro1 is a Ca^2+^-sensing adaptor protein that links mitochondria to the trafficking apparatus, for their microtubule-dependent transport along axons and dendrites, in order to meet the metabolic and Ca^2+^-buffering requirements of the cell. Here, we explore the role of Miro1 in parvalbumin interneurons and how changes in mitochondrial trafficking could alter brain network activity. By employing live and fixed imaging, we found that the impairments in Miro1-directed trafficking in PV+ interneurons altered their mitochondrial distribution and axonal arborization. These changes were accompanied by an increase in the *ex vivo* hippocampal γ-oscillation (30 – 80 Hz) frequency and promoted anxiolysis. Our findings show that precise regulation of mitochondrial dynamics in PV+ interneurons is crucial for proper neuronal signaling and network synchronization.

## INTRODUCTION

Parvalbumin (PV+) interneurons constitute a small proportion of the total neuronal population (less than 2% in the hippocampus) yet they possess crucial roles in shaping neuronal network activity (Freund and Buzsáki, 1996; Jonas et al., 2004; Pelkey et al., 2017). PV+ interneurons inhibit their postsynaptic targets efficiently by applying fast perisomatic inhibition, and have been directly implicated in the generation of network activity at the gamma (γ) band frequency (30 - 80 Hz) (Antonoudiou et al., 2020; Cardin et al., 2009; Hájos et al., 2004; Mann et al., 2005; Sohal et al., 2009). Network oscillations at γ-band frequency are believed to facilitate information transmission though circuit synchronization and local gain control that may be instrumental in multiple cognitive processes such as attention, learning and memory (Akam and Kullmann, 2010; Fries, 2015; Howard et al., 2003; Montgomery and Buzsáki, 2007; Sohal, 2016). Importantly, these oscillations are thought to be metabolically very costly and it has therefore been postulated that PV+ interneurons require substantial amounts of energy via ATP hydrolysis to sustain the high firing rate and dissipate ion gradients during neuronal transmission (Attwell and Laughlin, 2001; Kann, 2011, 2016; Kann and Kovács, 2007; Kann et al., 2014). Thus, it is crucial to understand the metabolic expenditure and the involvement of mitochondria in PV+ interneurons. Indeed, electron microscopy, histochemical and transcriptomic approaches have revealed that PV+ interneurons have a higher density of energy-producing mitochondria and elevated expression levels of electron transport chain components (Adams et al., 2015; Gulyás et al., 2006; Nie and Wong-Riley, 1995; Paul et al., 2017).

The spatiotemporal organization of mitochondria is essential for the precise provision of ATP and Ca^2+^-buffering for neuronal transmission and communication (Devine and Kittler, 2018; MacAskill and Kittler, 2010). Miro1 is a mitochondrial adaptor protein, responsible for coupling mitochondria to the cytoskeleton and for their bidirectional trafficking in axons and dendrites (Birsa et al., 2013; Guo et al., 2005; López-Doménech et al., 2016; López-Doménech et al., 2018; Macaskill et al., 2009; Nguyen et al., 2014; Saotome et al., 2008; Wang and Schwarz, 2009). Global deletion of Miro1 (encoded by the *Rhot1* gene) is perinatal lethal, while the conditional removal of Miro1 from cortical and hippocampal pyramidal cells alters the occupancy of dendritic mitochondria due to impairment in trafficking, resulting in dendritic degeneration and cell death (López-Doménech et al., 2016). In contrast, the significance of mitochondrial trafficking and distribution in PV+ interneurons, and the role of Miro1, is completely unexplored and especially interesting as their axon is highly branched with a cumulative length reaching up to 50 mm in the hippocampus (Hu et al., 2014).

In this study, we generated a transgenic mouse line where mitochondria are fluorescently labelled in PV+ interneurons. We crossed this line with the Miro1 floxed mouse (*Rhot1*^(fl/fl)^), to generate a model where Miro1 was conditionally knocked-out exclusively in PV+ interneurons. Using two-photon live-imaging of *ex vivo* organotypic brain slices, we demonstrated a reduction in mitochondrial trafficking in the absence of Miro1 in PV+ interneurons in the hippocampus. The impairment in Miro1-directed mitochondrial transport led to an accumulation of mitochondria in the soma and their depletion from axonal presynaptic terminals in acute hippocampal brain slices. Loss of Miro1 resulted in alterations in axonal but not dendritic branching in PV+ interneurons. This was accompanied by an increased frequency of γ-oscillations in hippocampal brain slices and a reduction in anxiety-related emotional behavior. Thus, we show that Miro1-dependent mitochondrial positioning is essential for correct PV+ interneuron function, network activity and anxiolytic animal behavior.

## RESULTS

### Loss of Miro1 in parvalbumin interneurons impairs mitochondrial trafficking

Mitochondrial enrichment in parvalbumin (PV+) interneurons is thought to reflect the high energetic demands of these cells (Gulyás et al., 2006). Consistent with this finding, we also observed that the protein levels of subunit IV of cytochrome c oxidase (COX-IV) were elevated in PV immuno-positive regions (3.6×10^−6^ ± 1.33×10^−5^ a.u.) when compared to immuno-negative areas in hippocampal slices (2.8×10^−6^ ± 1.26×10^−5^ a.u., **Fig 1A**, p < 0.0001), further supporting that PV+ interneurons heavily rely on mitochondria. Yet, mitochondrial dynamics in PV+ interneurons have not been explored. To specifically examine the role of mitochondrial transport in PV+ interneurons, we disrupted the mitochondrial adaptor protein Miro1 by crossing the PV^Cre^ mouse line with the *Rhot1^(^*^fl/fl)^ mouse (López-Doménech et al., 2016), thus generating a model where Miro1 was selectively knocked-out in PV+ interneurons (**Supplementary Fig 1A**). We then crossed this PV^Cre^ *Rhot1^(^*^fl/fl)^ line with a transgenic mouse where mitochondria expressed the genetically encoded Dendra2 fluorophore (MitoDendra) (Pham et al., 2012) (**Supplementary Fig 1A**), allowing for the visualization of mitochondria selectively in PV+ interneurons (**Supplementary Fig 1B**). Thus, we generated a mouse where Miro1 was knocked-out and mitochondria were fluorescently labelled selectively in PV+ interneurons (**Supplementary Fig 1C**). Gene expression under the PV+ promoter begins around P10 and is stabilized around P28 (Barnes et al., 2015). Thus, Cre-dependent removal of Miro1 is not expected to impact on the migration, differentiation and viability of PV+ neurons (Okaty et al., 2009; del Río et al., 1994). The Miro1 fluorescence intensity was significantly reduced in brain slices from the knock-out mouse, confirming the selective removal of Miro1 from PV+ interneurons (Miro^(+/Δ)^ 844 ± 32.8 a.u., Miro^(Δ/Δ)^ 518 ± 20.5 a.u. **Supplementary Fig 1D**, p < 0.0001). These immunohistochemical experiments demonstrate that the PV^Cre^ *Rhot1^(^*^fl/fl)^ MitoDendra mouse can be utilized to examine mitochondria dynamics in parvalbumin interneurons in a system where Miro1 is absent.

**Figure 1.**
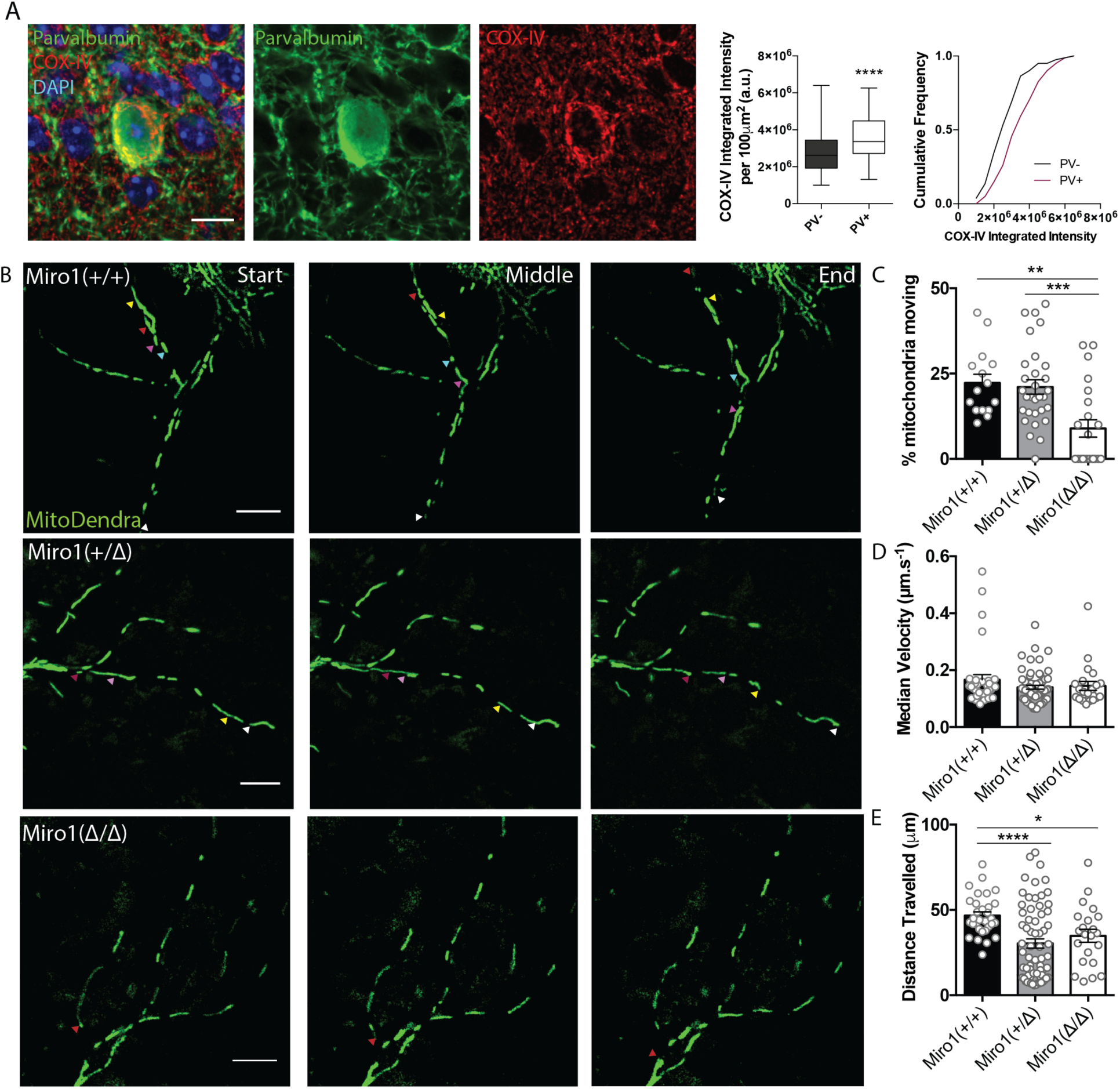
Cell-type specific removal of Miro1 from parvalbumin interneurons impairs mitochondrial trafficking. **A.** COX-IV levels in PV+ interneurons. The fluorescence signal of COX-IV is increased in PV immuno-positive cells in the hippocampus. Scale Bar = 10 μm. Boxplot and cumulative distribution show the quantification of the mean integrated intensity of the COX-IV fluorescent signal in PV immuno-positive and -negative regions (n = 81 cells, 11 slices, 3 animals) **B.** Miro1-directed mitochondrial trafficking in organotypic brain slices. Representative images from a 500 second two-photon movie of MitoDendra+ mitochondria in PV+ interneurons in *ex vivo* organotypic hippocampal slices from Miro1^(+/+)^, Miro1^(+/Δ)^ and Miro1^(Δ/Δ)^ animals. The colored arrows denote the position of individual mitochondria during the movie. Scale bar = 10 μm **C.** Quantification of the percentage of moving mitochondria (n_Miro(+/+)_ = 15 movies, 9 slices, 4 animals, n_Miro(+/Δ)_ = 32 movies, 10 slices, 4 animals, n_Miro(Δ/Δ)_ = 22 movies, 8 slices, 4 animals) **D.** Quantification of the median velocity (n_Miro(+/+)_ = 35 mitochondria, n_Miro(+/Δ)_ = 65 mitochondria, n_Miro(Δ/Δ)_ = 22 mitochondria) **E.** Quantification of the distance travelled (n_Miro(+/+)_ = 35 mitochondria, n_Miro(+/Δ)_ = 65 mitochondria, n_Miro(Δ/Δ)_ = 22 mitochondria).

Next, we wanted to investigate the contribution of Miro1 to mitochondrial trafficking in PV+ interneurons. We therefore performed two-photon live-imaging in intact organotypic brain tissue (at 7-9 days *in vitro*) from neonatal (P6-8) control (Miro1^(+/+)^), hemi-floxed (Miro1^(+/Δ)^) and conditional knock-out (Miro1^(Δ/Δ)^) animals (**Fig 1B**). Consistent with other models where Miro1 is knocked-out (López-Doménech et al., 2016; Nguyen et al., 2014), we found a significant reduction in the percentage of moving mitochondria in the absence of Miro1 when compared to the control (Miro1^(+/+)^ 23 ± 3%, Miro1^(+/Δ)^ 21 ± 2%, Miro1^(Δ/Δ)^ 9 ± 3%, **Fig 1C**, p_Miro1(+/Δ)_ = 0.939, p_Miro1(Δ/Δ)_ = 0.002). Even though the moving mitochondria exhibited similar velocities in the three conditions (Miro1^(+/+)^ 0.17 ± 0.018 μm/s, Miro1^(+/Δ)^ 0.14 ± 0.007 μm/s, Miro1^(Δ/Δ)^ 0.15 ± 0.016 μm/s, **Fig 1D**, p_Miro1(+/Δ)_ = 0.229, p_Miro1(Δ/Δ)_ = 0.557), the length of the travelled trajectory was significantly shorter in Miro1^(+/Δ)^ (31 ± 2.7 μm) and Miro1^(Δ/Δ)^ (35 ± 3.8 μm) when compared to the control (49 ± 2.0 μm, **Fig 1E**, p_Miro1(+/Δ)_ < 0.0001, p_Miro1(Δ/Δ)_ = 0.024). This experiment demonstrates that the presence of Miro1 is critical for mitochondrial trafficking in PV+ interneurons.

### Loss of Miro1 in parvalbumin interneurons results in mitochondrial accumulation in the soma and depletion from axonal presynaptic terminals

Given that Miro1 is a crucial component of the mitochondrial transport machinery in PV+ cells, we wanted to investigate the effect of impaired trafficking on mitochondrial localization in PV+ interneurons, *ex vivo.* We noticed that mitochondria accumulated in the cell bodies of PV+ interneurons depleted of Miro1 in hippocampal sections from adult mice (**Fig 2A**). The number of PV immuno-positive cells that contained mitochondrial accumulations in the soma was significantly increased when Miro1 was knocked-out (58 ± 5.7%), compared to control (11 ± 3.1%, **Fig 2B**, p <0.0001). This is consistent with the somatic mitochondrial accumulations that have also been reported in pyramidal cells in the CaMK-II^Cre^ *Rhot1^(^*^fl/fl)^ model (López-Doménech et al., 2016) and are presumably due to the inefficient trafficking of mitochondria out to the neurites of the cell. The mitochondria in the Miro1^(Δ/Δ)^ occupied a smaller area relative to the cell body (Miro1^(+/Δ)^ 39 ± 1.0%, Miro1^(Δ/Δ)^ 28 ± 1.0%) and seem to be concentrated around the nucleus when Miro1 was knocked-out (**Fig 2C**, p < 0.0001). Together, these data suggest that the spatial distribution of mitochondria is altered in PV+ interneurons in the absence of Miro1.

**Figure 2.**
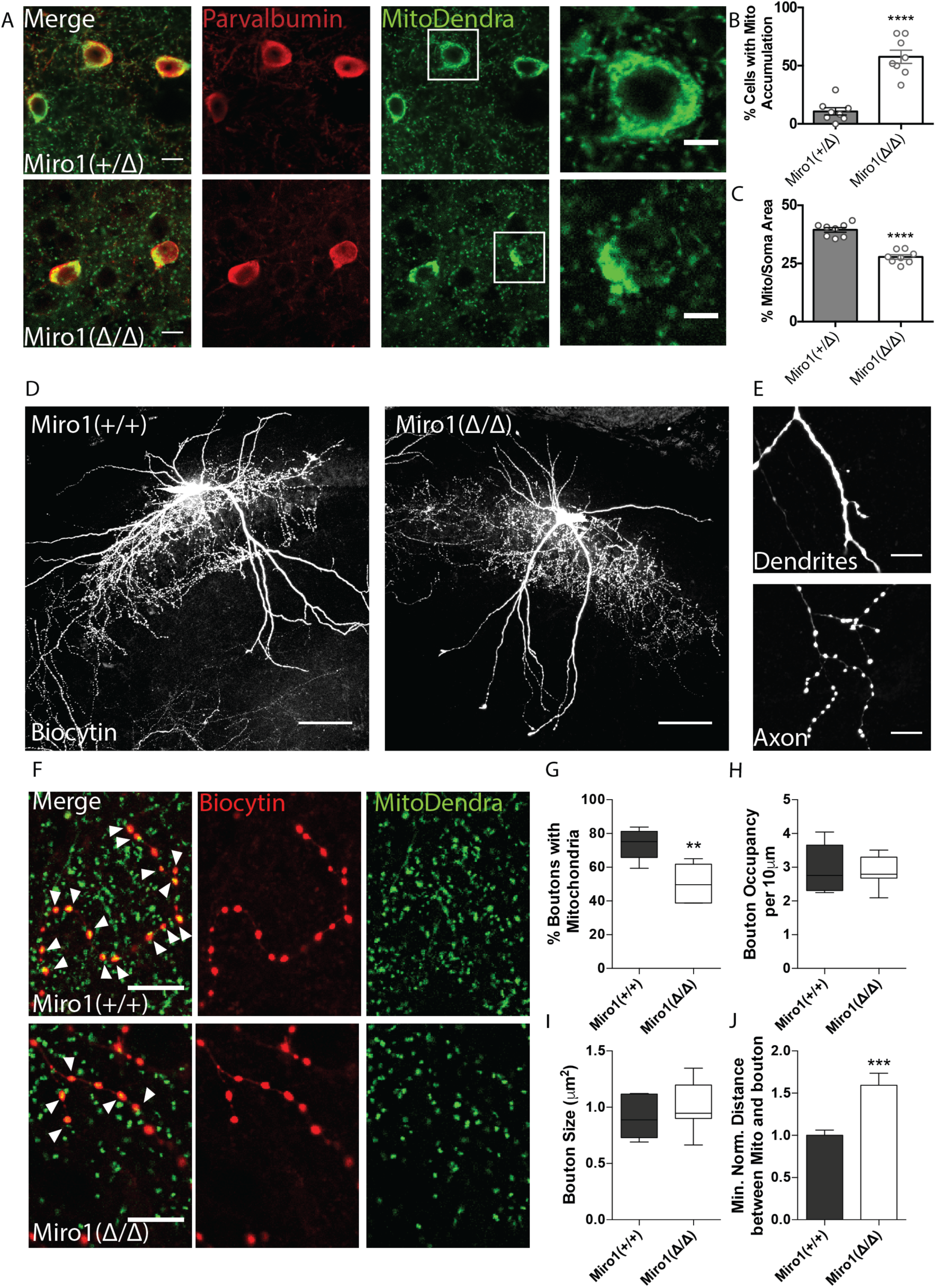
Loss of Miro1 results in an accumulation of mitochondria in the soma and depletion from axonal presynaptic terminals. **A.** Loss of Miro1 results in MitoDendra+ accumulation in the somata of PV+ interneurons. Confocal images from Miro1^(+/Δ)^ and Miro1^(Δ/Δ)^ PV immuno-positive cells in fixed brain tissue. Scale bar = 10 μm. Cells in the white box are zoomed as illustrated in the images on the right, to clearly depict the mitochondrial accumulations in the Miro1^(Δ/Δ)^. Scale bar = 5 μm. **B.** Bar chart shows the quantification for the percentage of cells that contain mitochondrial clusters (n_Miro(+/Δ)_ = 8 slices, 3 animals and n_Miro(Δ/Δ)_ = 8 slices, 3 animals) **C.** Quantification for the percentage area that mitochondria occupy within the PV immuno-positive soma. Due to the presence of mitochondrial accumulations the %Mito/Soma area is significantly decreased in the Miro1^(Δ/Δ)^ (n_Miro(+/Δ)_ = 8 slices, 3 animals and n_Miro(Δ/Δ)_ = 8 slices, 3 animals). **D.** Representative max-projected confocal stack of biocytin-filled PV+ interneurons in the hippocampus of 350 μm fixed acute brain slices. Scale bar = 100 μm **E.** Example of the distinct distribution of biocytin in dendritic and axonal compartments. Biocytin is diffused in dendrites (top) and clustered in axonal terminals (bottom) **F.** Loss of Miro1 depletes mitochondria from axonal presynaptic terminals. Representative images of biocytin-filled synaptic boutons (red) and mitochondria (green) in Miro1^(+/+)^ and Miro1^(Δ/Δ)^ neurons. Arrows point to boutons that contain mitochondria. Scale bar = 5 μm **G.** Boxplots for the quantification of the percentage of boutons that contain mitochondria (n_Miro(+/+)_ = 5 neurons, 4 slices, 2 animals and n_Miro(Δ/Δ)_ = 7 neurons, 6 slices, 3 animals) **H.** Boxplots for the quantification of the occupancy of boutons in axonal segments (n_Miro(+/+)_ = 5 neurons, 4 slices, 2 animals and n_Miro(Δ/Δ)_ = 7 neurons, 6 slices, 3 animals) **I.** Quantification of the mean size of boutons (n_Miro(+/+)_ = 625 boutons, 5 neurons, 4 slices, 2 animals, n_Miro(Δ/Δ)_ = 526 boutons, 7 neurons, 6 slices, 3 animals) **J.** Quantification for the minimum normalised distance between boutons and mitochondria (n_Miro(+/+)_ = 625 boutons, 5 neurons, 4 slices, 2 animals, n_Miro(Δ/Δ)_ = 526 boutons, 7 neurons, 6 slices, 3 animals).

To further understand the impact of impaired Miro1-dependent mitochondrial trafficking in PV+ interneurons, we looked at the organization of mitochondria in the axons and dendrites of PV+ cells. MitoDendra+ cells were biocytin-filled allowing for the morphological visualization of individual PV+ interneurons in acute brain slices (**Fig 2D**). Biocytin was uniformly diffused along dendrites and accumulated in the axon allowing for the identification of distinct presynaptic boutons (**Fig 2E, 2F**) (Swietek et al., 2016). We observed that 74 ± 4.1% of presynaptic terminals contained MitoDendra+ mitochondria in control cells (**Fig 2F, 2G**). This is consistent with literature where approximately 75% of PV+ axonal boutons are enriched with mitochondria, and this is in contrast to pyramidal cells where less than half are associated with mitochondria (Glausier et al., 2017; Kwon et al., 2016; Smith et al., 2016; Vaccaro et al., 2017). However, we report a marked reduction in the axonal presynaptic terminals that contained mitochondria in the cells where Miro1 was knocked-out (51 ± 4.0%, **Fig 2G**, p = 0.003). The distribution along the axon (occupancy) and size of boutons were unaffected by the loss of Miro1 (**Fig 2H, 2I**). The minimum normalised distance between boutons and mitochondria was increased in Miro1^(Δ/Δ)^ PV+ interneurons (1.6 ± 0.1 a.u.) when compared to Miro1^(+/+)^ (1 ± 0.1 a.u.) suggesting that mitochondria might no longer be captured in the presynapse as they are found further away from the axonal terminals (**Fig 2J**, p = 0.0001). These data indicate that defects in Miro1-directed mitochondrial trafficking are sufficient to shift the mitochondrial distribution along the axon, leading to a depletion of mitochondria away from axonal presynaptic boutons, *in vivo*.

### Loss of Miro1 in PV+ interneurons results in an enrichment of mitochondria close to axonal branching sites and an increase in axonal branching

Our data suggest that the impairment in Miro1-dependent trafficking altered the precise mitochondrial positioning in PV+ interneuron axons. To examine whether this alteration could result in a structural change in PV+ interneuron morphology due to a reorganization of the mitochondrial network in the absence of Miro1, we used the biocytin fill to generate a neuronal reconstruction (**Fig 3A**). The total length of PV+ interneuron dendrite and axon did not differ between control and conditional knock-out cells (**Fig 3B**). We noticed an increase in the number of processes in Miro1^(Δ/Δ)^ cells (506 ± 34 processes) when compared to Miro1^(+/+)^ (376 ± 46 processes, **Fig 3C**, p = 0.042) and a trend towards an increase in the number of branches between Miro1^(+/+)^ (342 ± 42 branches) and Miro1^(Δ/Δ)^ conditions (465 ± 35 branches, **Fig 3D**, p = 0.052). We further investigated whether the increase in processes was attributed to a selective increase in the branches of axons or dendrites in the absence of Miro1. By visually assessing and independently labelling axons and dendrites during the reconstruction process, we isolated the contribution of each compartment to the total number of branches. Interestingly, there was a selective increase in axonal (**Fig 3F, 3H**) but not dendritic branching (**Fig 3E, 3G**) in Miro1^(Δ/Δ)^ (427 ± 32 branches) when compared to Miro1^(+/+)^ (296 ± 38 branches, **Fig 3F**, p = 0.031). By performing Sholl analysis and plotting the number of intersections as a function of the distance from the soma in Miro1^(+/+)^ and Miro1^(Δ/Δ)^ cells, we noticed an enhancement in axonal branching in the proximal axon (between 50 μm and 200 μm from the cell body) when Miro1 was knocked-out (**Fig 3H**). By using the neuronal reconstruction as a mask, we isolated the MitoDendra+ mitochondrial network in individual PV+ interneurons and compared the mitochondrial organization in control and knock-out cells. Sholl analysis revealed a shift in mitochondrial distribution proximal to the soma in Miro1^(Δ/Δ)^ cells (**Fig 3J**). We quantified the minimum normalised distance between branch points and mitochondria (**Fig 3I**) in Miro1^(+/+)^ (1 ± 0.06 a.u.) and Miro1^(Δ/Δ)^ (0.7 ± 0.04 a.u.) and found a decrease in the absence of Miro1 (**Fig 3K**, p = 0.0003). Additionally, the probability of encountering a mitochondrion within 1 μm from the branch point is higher in the Miro1^(Δ/Δ)^ (P = 0.49) when compared to the control condition (P = 0.31) (data not shown). Thus, the loss of Miro1-directed trafficking not only shifted mitochondria away from the presynapse but also placed them closer to locations of axonal branching. The close proximity of mitochondria to axonal branching points in the absence of Miro1 suggests roles for their involvement in driving the formation of axonal segments.

**Figure 3.**
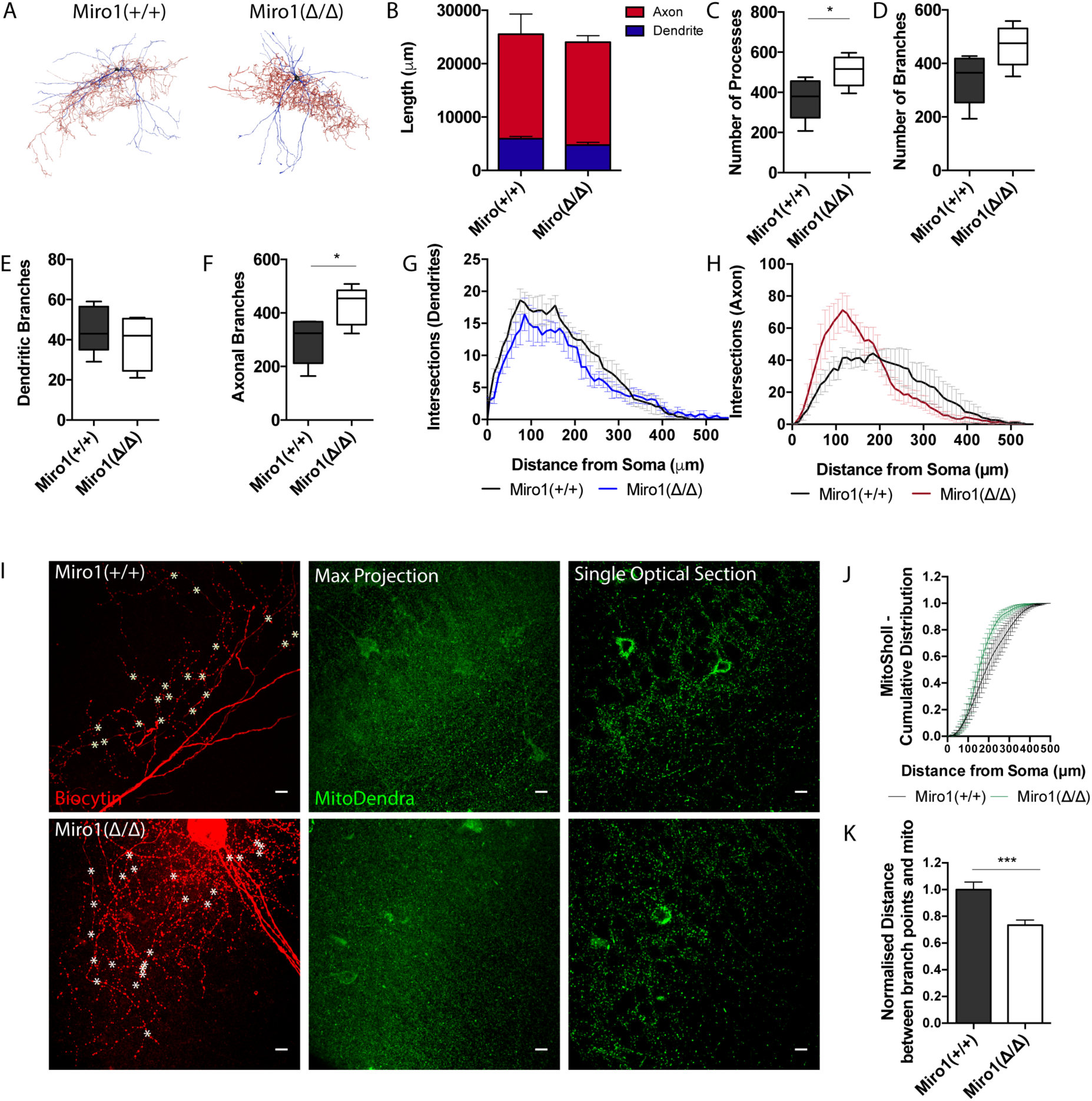
Loss of Miro1 results in increased axonal branching in hippocampal parvalbumin interneurons. **A.** Max-projected reconstruction of the neurons in Fig 2D from Neuromantic software. Dendrites are depicted in blue and axons in red. **B.** Quantification of the neuronal length with the dendritic and axonal contribution depicted in blue and red respectively. **C.** Boxplot for the quantification of the total number of processes **D.** Boxplot for the quantification of the total number of branches **E.** Box plot for the quantification of the number of dendritic branches. **F.** Box plot for the quantification of the number of axonal branches. **G.** Number of intersections between dendritic branches and Sholl rings are plotted at distances away from the soma **H.** Number of intersections between axonal branches and Sholl rings are plotted at distances away from the soma. (n_Miro(+/+)_ = 5 neurons, 3 slices, 2 animals, n_Miro(Δ/Δ)_ = 5 neurons, 4 slices, 3 animals). **I.** Loss of Miro1 results in mitochondria being found closer to points of axonal branching. High magnification (63x) max-projected confocal stacks of biocytin in a 350 μm hippocampal slice. Analysis was performed in 3D in single optical sections. White stars denote branch points. Scale bar = 10 μm. **J.** Cumulative distribution of the MitoDendra+ mitochondrial network (MitoSholl) in individual PV+ interneuron (n_Miro(+/+)_ = 5 neurons, 3 slices, 2 animals, n_Miro(Δ/Δ)_ = 4 neurons, 4 slices, 3 animals). **K.** Bar graph shows the normalised minimum distance between a branch point and a mitochondrion in the confocal stack (n_Miro(+/+)_ = 122 branch points, 5 neurons, 4 slices, 2 animals and n_Miro(Δ/Δ)_ = 171 branch points, 7 neurons, 6 slices, 3 animals).

### Loss of Miro1-dependent mitochondrial positioning does not alter parvalbumin interneuron mediated inhibition

We next wanted to address whether the alterations in mitochondrial localisation due to impaired trafficking could affect parvalbumin interneuron function and their ability to apply fast perisomatic inhibition in the absence of Miro1. To do that, we recorded the spontaneous inhibitory postsynaptic currents (sIPSCs) that pyramidal cells received in the hippocampus in acute brain slices from Miro1^(+/+)^ and Miro1^(Δ/Δ)^ animals (**Supplementary Fig 2A**). Neither the frequency of the sIPSCs, represented as the inter event interval, nor the amplitude of the responses were different between the two conditions (**Supplementary Fig 2B, 2C**), suggesting that the loss of Miro1 did not lead to an alteration in inhibitory synaptic transmission. The recorded sIPSCs however could contain the inhibitory contribution of other interneurons in the network. To specifically assess the PV+ interneuron mediated inhibition, we generated a mouse model where PV+ interneurons could be temporally controlled by the light-induced activation (photoactivation) of channel rhodopsin 2 (ChR2) (**Supplementary Fig 2D, 2E**). We then photoactivated PV+ interneurons (1ms pulse width, 30 repetitions per cell) and quantified the mean evoked IPSCs (eIPSCs) neighbouring pyramidal cells received in the hippocampus (**Supplementary Fig 2E, 2F)**. We found that there was no change in the amplitude (**Supplementary Fig 2G**), charge transfer (**Supplementary Fig 2H**), and decay (**Supplementary Fig 2I**) between control and Miro1^(Δ/Δ)^ conditions. These data strongly suggest that the PV+ interneuron ability to apply perisomatic inhibition is unaffected by the loss of Miro1 and subsequent changes in mitochondrial dynamics. We also tested whether PV+ cells could sustain inhibition and recover after long-lasting light-activation (**Supplementary Fig 2J**). We presented a 2 second light train stimulation (40 Hz; 1ms pulse width) that was repeated 10 times and recorded the eIPSCs from neighbouring pyramidal cells. In order to assess whether the cells recovered in a similar manner after the photo-stimulation, we presented one light pulse at increasing time intervals at the end of every train and measured the eIPSC response. There was no difference in neither the amplitude of the inhibitory currents received by pyramidal cells during the 2 second stimulation (**Supplementary Fig 2K**) nor in the recovery of post-light train pulses. Thus, Miro1-dependent mitochondrial positioning does not seem to alter the synaptic properties of PV+ cells and may not be important for short term recovery of the inhibitory responses either, as control and knock-out cells behave similarly (**Supplementary Fig 2L**). In conclusion, the loss of Miro1 does not seem to alter inhibitory properties of PV+ interneurons in the hippocampus of two month old animals.

### Parvalbumin interneurons receive increased excitatory inputs when Miro1 is conditionally knocked-out

Next, we sought to understand whether the loss of Miro1 and subsequent changes in mitochondrial dynamics alter the intrinsic features of PV+ interneurons. We recorded the intrinsic properties of PV+ interneurons in Miro1^(+/+)^, Miro1^(+/Δ)^ and Miro1^(Δ/Δ)^ slices by applying depolarising and hyperpolarising pulses (**Supplementary Fig 3A**). We observed no difference in action potential (AP) peak amplitude (**Supplementary Fig 3B**), input resistance (**Supplementary Fig 3E**), threshold to fire (**Supplementary Fig 3F**), and spike rate at 40 pA above rheobase current (**Supplementary Fig 3D**) in the three conditions. The AP half-width was slightly higher in Miro1^(+/+)^ (0.4 ± 0.03 ms) when compared to Miro1^(+/Δ)^ (0.3 ± 0.02 ms) and Miro1^(Δ/Δ)^ (0.3 ± 0.02 ms) recordings (**Supplementary Fig 3C**, p_Miro1(+/Δ)_ = 0.048, p_Miro1(Δ/Δ)_ = 0.864). The membrane time constant was slightly decreased in Miro1^(+/Δ)^ (8 ± 0.8 ms) and Miro1^(Δ/Δ)^ (8 ± 0.6 ms) animals when compared to control (12 ± 2.0 ms, **Supplementary Fig 3G**, p_Miro1(+/Δ)_ = 0.073, p_Miro1(Δ/Δ)_ = 0.046). We speculate that although the differences in intrinsic properties were small, the loss of Miro1 from PV+ interneurons might potentially result in shorter integration time and faster response to excitatory inputs. The change in membrane time constant could also potentially reflect the changes to the morphology of the cell (Isokawa, 1997).

In order to investigate whether PV+ interneurons received differential synaptic inputs, we performed whole-cell recordings and acquired the spontaneous excitatory postsynaptic currents (sEPSCs) in the hippocampus of acute brain slices (**Fig 4G**). The sEPSC frequency was significantly increased, as shown by the decrease in the mean inter-event interval (IEI) between Miro1^(+/+)^ (0.03 ± 0.009 s) and Miro1^(Δ/Δ)^ (0.02 ± 0.003 s) cells (**Fig 4H**, p = 0.025). Additionally, the amplitude of sEPSCs was elevated in Miro1^(Δ/Δ)^ (−188 ± 17.2 pA) slices when compared to control slices (−130 ± 7.8 pA, **Fig 4I**, p = 0.0097). Thus, the increase in sEPSC frequency and amplitude suggest that PV+ interneurons receive an enhanced excitatory drive when Miro1 was knocked-out.

**Figure 4.**
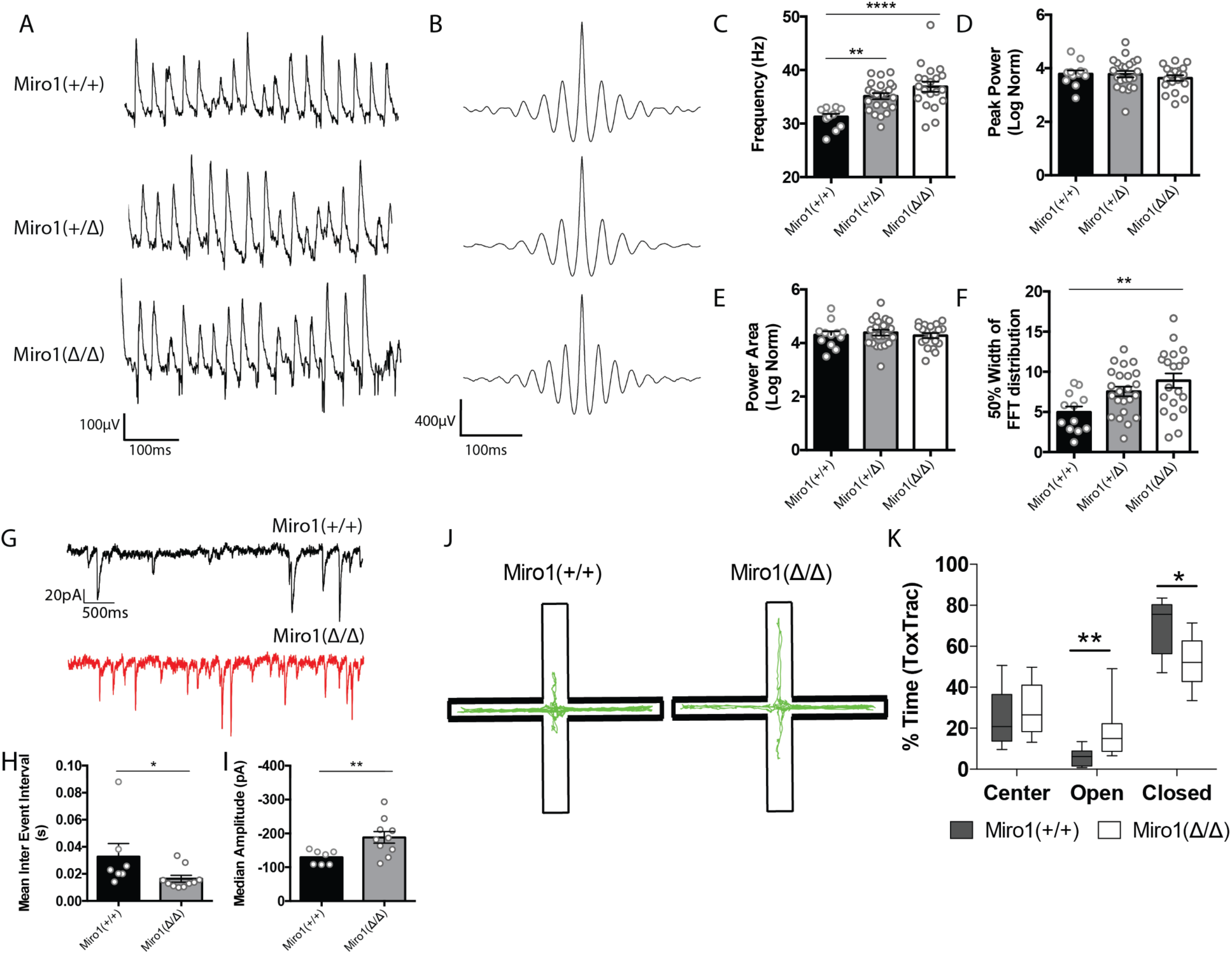
Miro1 knock-out results in altered hippocampal network activity and anxiety-related behavior. **A.** Loss of Miro1 increases the frequency of γ-oscillations. Representative local field potential recordings from the stratum pyramidale of the CA3 hippocampal area in acute brain slices from Miro1^(+/+)^, Miro1^(+/Δ)^ and Miro1^(Δ/Δ)^ slices **B.** Representative auto-correlogram of γ-oscillations from Miro1^(+/+)^, Miro1^(+/Δ)^ and Miro1^(Δ/Δ)^ animals **C.** Quantification of the peak frequency **D.** Quantification of the normalised peak power. **E**. Quantification of the normalised power area **F.** Quantification of the 50% width dispersion of the FFT distribution (n_Miro(+/+)_ = 12 slices, 2 animals, n_Miro(+/Δ)_ = 23 slices, 6 animals and n_Miro(Δ/Δ)_ = 20 slices, 6 animals). **G.** Miro1^(Δ/Δ)^ PV+ interneurons received increased glutamateric input. Representative electrophysiological traces from Miro1^(+/+)^ (black) and Miro1^(Δ/Δ)^ (red) cells. **H.** Quantification for the mean inter-event interval (IEI). **I.** Quantification for the median amplitude (n_Miro(+/+)_ = 7 recordings, 2 animals and n_Miro(Δ/Δ)_ = 10 recordings, 2 animals). **J.** Assessment of anxiety-related behavior using the elevated plus maze (EPM). Schematic diagram of the EPM and representative ToxTrac trajectories (green) from Miro1^(+/+)^ and Miro1^(Δ/Δ)^ animals. **K.** Boxplot for the quantification of the percentage of time that Miro1^(+/+)^ and Miro1^(Δ/Δ)^ animals spent in the closed, open arms and center of the EPM (n_Miro(+/+)_ = 9 animals, n_Miro(Δ/Δ)_ = 8 animals)

### Alterations in Miro1-dependent mitochondrial positioning change hippocampal network activity and anxiety-related animal behavior

We then sought to examine whether the Miro1-dependent alterations in mitochondrial distribution could result in altered neuronal network activity. Given that γ-oscillations are energetically very costly (Kann et al., 2014) and depend on proper PV+ interneuron function, we wanted to see whether the loss of Miro1 could have an impact on the ability of PV+ cells to synchronize neuronal networks. We measured carbachol induced γ-oscillations in the CA3 area of the hippocampus by recording local field potentials in acute brain slices of Miro1^(+/+)^, Miro1^(+/Δ)^ and Miro1^(Δ/Δ)^ animals (**Fig 4A, 4B**). Even though the peak power and power area were not significantly different (**Fig 4D, 4E**), we found a small but significant increase in the peak frequency of the oscillations that appeared to be gene-dose dependent between the different genotypes (Miro1^(+/+)^ 31 ± 0.6 Hz, Miro1^(+/Δ)^ 35 ± 0.6 Hz, Miro1^(Δ/Δ)^ 37 ± 0.9 Hz, **Fig 4C**, p_Miro1(+/Δ)_ = 0.004, p_Miro1(Δ/Δ)_ < 0.0001). We then quantified the 50% width of the power spectrum distribution as a measure of the oscillation variability (**Fig 4F**). We noticed that the distribution was significantly broader in Miro1^(+/Δ)^ (8 ± 0.6) and Miro1^(Δ/Δ)^ (9 ± 0.9) when compared to control (5 ± 0.7), suggesting increased variability of the γ-cycle duration (**Fig 4F**, p_Miro1(Δ/Δ)_ = 0.005). These data demonstrate that the loss of Miro1 and subsequent changes in mitochondrial trafficking and distribution in PV+ interneurons are sufficient to alter hippocampal network activity, further involving mitochondria in the modulation of *ex vivo* γ-oscillations (Inan et al., 2016; Kann, 2011; Kann et al., 2011).

Finally, we wanted to see whether the loss of Miro1 from PV+ interneurons was sufficient to induce functional changes in the behavior of these animals. We observed no changes in husbandry behavior (**Supplementary Fig 4A**) assessed by the shredding of Nestlets (Deacon, 2006), motor coordination and learning (**Supplementary Fig 4B**) on the rotarod (Deacon, 2013) and short-term memory (**Supplementary Fig 4C**) in the T-Maze (Deacon and Rawlins, 2006). We also did not observe a robust difference in spatial exploration of Miro1^(+/+)^ and Miro1^(Δ/Δ)^ animals in the open field assessment (**Supplementary Fig 4D**) (Seibenhener and Wooten, 2015). The velocity (**Supplementary Fig 4E**), distance travelled (**Supplementary Fig 4F**), and time spent in the areas of the arena (**Supplementary Fig 4G**) were not statistically different between the two genotypes. Finally, we tested anxiety-like behavior of littermate control and conditional knock-out animals in the elevated plus maze (EPM) (**Fig 4J**). As expected, both Miro1^(+/+)^ and Miro1^(Δ/Δ)^ mice spent the majority of time in the closed arms due to the natural aversive behavior to light. The Miro1^(Δ/Δ)^ animals however spent more time in the open arms (18 ± 5%) than their littermate controls (6 ± 1%, **Fig 4K**, p = 0.004). The performance of Miro1^(Δ/Δ)^ animals in the EPM suggests that they may exhibit a reduced anxiety-like phenotype. In conclusion, the selective loss of Miro1 from parvalbumin interneurons results in changes in mitochondrial dynamics, axonal remodeling and network activity that might give rise to anxiety related phenotypes, affecting the behavior of these animals.

## DISCUSSION

In this study, we demonstrate that the conditional removal of Miro1 in PV+ cells resulted in a mitochondrial trafficking impairment. Furthermore, the loss of Miro1 led to an accumulation of mitochondria in the soma of PV+ cells and their depletion from axonal presynaptic terminals. The relocation of mitochondria closer to sites of axonal branching was associated with a selective enrichment in axonal branches proximal to the cell body. Interestingly, these animals also exhibited faster γ-oscillations and a reduced anxiety-like phenotype. Our data suggest that Miro1-dependent mitochondrial positioning is implicated in shaping hippocampal network activity and animal behavior.

The enrichment of mitochondria in PV+ interneurons denotes their important role in the correct function of these neurons (Gulyás et al., 2006; Inan et al., 2016; Lin-Hendel et al., 2016; Paul et al., 2017). Our immunohistochemical data also demonstrate that PV+ interneurons display increased levels of the subunit IV of cytochrome c oxidase (COX-IV) (**Fig 1A**). This is consistent with evidence proposing high levels of oxidative phosphorylation proteins such as complex I, cytochrome C oxidase, cytochrome C and ATP synthase in these cells (Gulyás et al., 2006; Kann et al., 2011; Paul et al., 2017). This observation could further support the notion that PV+ interneurons are metabolically more active to meet their high energy demands and suggests that stringent dependence on mitochondrial function could render PV+ interneurons susceptible to incidents of mitochondrial damage.

The generation of the PV^Cre^ *Rhot1*^(fl/fl)^ MitoDendra transgenic mouse permitted the visualization of mitochondria and the conditional removal of Miro1 specifically in PV+ cells (**Supplementary Fig 1**). Given the role of the adaptor protein Miro1 in the bidirectional trafficking of mitochondria to locations of high energy demand (Birsa et al., 2013; Guo et al., 2005; López-Doménech et al., 2016; López-Doménech et al., 2018; Macaskill et al., 2009; Nguyen et al., 2014; Saotome et al., 2008; Wang and Schwarz, 2009), we examined the importance of correct mitochondrial transport in PV+ interneurons. The conditional removal of Miro1 resulted in a decrease in mitochondrial trafficking, consistent with the functional role of Miro1 (**Fig 1B**). A small subset of mitochondria remained mobile in the absence of Miro1, indicating that mitochondrial transport was not completely abolished in Miro1^(Δ/Δ)^. Other studies also report a reduction, but not complete cessation of mitochondrial transport, upon Miro1 loss (López-Doménech et al., 2016; Macaskill et al., 2009; Russo et al., 2009), hinting to the existence of alternative mitochondrial transport mechanisms. The physical attachment of mitochondria to other motor-adaptor proteins might still facilitate organelle transport. Our lab has demonstrated that Trak1/2 motors can still be recruited to the mitochondria in mouse embryonic fibroblasts (MEFs), in a Miro1-independent fashion (López-Doménech et al., 2018). The mobile mitochondria in PV+ interneurons exhibited shorter travelled trajectories in Miro1^(Δ/Δ)^ (**Fig 1E**). Thus, it is also possible that the remaining moving mitochondria engaged with cytoskeletal elements other than microtubules. Indeed, involvement of the actin cytoskeleton has been implicated in short-range mitochondrial movement (Chada and Hollenbeck, 2004; Morris and Hollenbeck, 1995). Still, the remaining moving mitochondria in our model exhibited directional, rather than randon movement, which favors transport along microtubules.

The loss of Miro1 from PV+ interneurons resulted in a mitochondrial accumulation in the cell bodies suggesting that mitochondria are no longer able to exit the soma due to impaired mitochondrial trafficking (**Fig 2A**). Similar perinuclear clustering has also been reported upon the loss of Miro1 in MEFs (López-Doménech et al., 2018; Nguyen et al., 2014) and in the CaMK-II^Cre^ *Rhot1*^(fl/fl)^ mouse at four months of age (López-Doménech et al., 2016). The loss of Miro1-directed mitochondrial transport was accompanied by a redistribution in the mitochondrial network in PV+ interneurons. While the majority of PV+ axonal boutons contain mitochondria (**Fig 2G**), loss of Miro1 resulted in their depletion from these sites (**Fig 2F**), suggesting a role for Miro1 in mitochondrial capture in presynaptic terminals along the PV+ interneuron axon. Mutations in dMiro, the drosophila orthologue, impair mitochondrial trafficking and deplete mitochondria from the axon, altering the morphology of both the axon and synaptic boutons in the neuromuscular junction (Guo et al., 2005). Miro1 also mediates the activity-dependent repositioning of mitochondria to synapses in excitatory neurons (Vaccaro et al., 2017). Collectively, this evidence proposes a role for Miro1-directed mitochondrial trafficking in the fine tuning of presynaptic mitochondrial occupancy. We speculate that the loss of mitochondria from axonal terminals in Miro1^(Δ/Δ)^ could be the consequence of defective protein-protein interactions with the tethering machinery, resulting in the inability of mitochondria to be retained in the presynaptic space. Mitochondrial transport on actin filaments mediates local distribution along the axon and destabilisation of F-actin reduces mitochondrial docking (Chada and Hollenbeck, 2003, 2004; Hirokawa et al., 2010; Shlevkov et al., 2019; Smith and Gallo, 2018). It is therefore possible that the loss of Miro1 might have disrupted the interaction between mitochondria and the actin cytoskeleton resulting in reduced presynaptic capture for example due to impairments in the Myo19-dependent coupling to actin in the absence of Miro1 (López-Doménech et al., 2018).

Neuronal mitochondrial misplacement is thought to impact on neuronal morphology as mitochondria are no longer precisely located in places where they are actively required to provide energy and buffer calcium (Devine and Kittler, 2018; Guo et al., 2005; Liu and Shio, 2008; López-Doménech et al., 2016). By reconstructing the morphology of PV+ interneurons in brain slices (**Fig 3A**), we reported a selective enrichment in axonal but not dendritic branching, proximal to the cell soma upon loss of Miro1 (**Fig 3H**). Interestingly, the loss of Miro1-directed mitochondrial trafficking and depletion of mitochondria from axonal boutons were accompanied by a general shift in the mitochondrial distribution closer to the soma (**Fig 3J**). Since mitochondria were found closer to sites of branch points in the absence of Miro1, our data suggest that the enhancement in axonal branching could be attributed to the perturbed mitochondrial distribution (**Fig 3I**). Mitochondrial arrest and presynaptic capture is necessary for axon extension and branching formation, as mitochondria can locally provide energy (Courchet et al., 2013; Sainath et al., 2017; Smith and Gallo, 2018; Spillane et al., 2013). The extension and retraction of the axon are developmentally dynamic processes (Chattopadhyaya et al., 2004, 2007; Huang et al., 2007) and the Miro1-dependent changes in mitochondrial distribution could promote either the formation of branches close to the cell body and/or the reorganization of existing branches.

We then examined whether the spatiotemporal regulation of mitochondrial positioning could impact on PV+ interneuron function and network activity. PV+ interneurons are critical elements in generating and maintaining γ-oscillations (Antonoudiou et al., 2020; Bartos et al., 2007; Buzsáki and Wang, 2012; Cardin et al., 2009; Sohal et al., 2009) and these oscillations heavily rely on intact mitochondrial function (Galow et al., 2014; Huchzermeyer et al., 2008, 2013, Kann et al., 2011, 2016; Whittaker et al., 2011). Indeed, pharmacological inhibition of complex I with rotenone, and uncoupling of oxidative phosphorylation with FCCP abolishes the power of γ-oscillations (Whittaker et al., 2011) while rotenone also reduced oxygen consumption in the CA3 area of the hippocampus, coupling respiration to network activity (Kann et al., 2011). By recording carbachol-induced rhythmic network activity in the CA3 area of the hippocampus, we reported a mild increase in the frequency and variability of γ-oscillations in a Miro1 dose dependent manner (**Fig 4C**). These results suggest that in addition to the proper functioning of mitochondria, their precise positioning along the axon might be important in the fine tuning of rhythmic oscillations in γ-frequency band. The width of the power spectra was wider when Miro1 was knocked-out from PV+ interneurons (**Fig 4F**), suggesting higher variability in the duration of γ-cycles. Alterations in axonal branching could have an implication on the innervation of postsynaptic targets (Huang et al., 2007) and it is therefore possible that the precise PV+ interneuron innervation of pyramidal cells changed (**Fig 3H**), resulting in an increase in the excitation PV+ interneurons received in the hippocampus (**Fig 4G, H**). We speculate that the extent of the axon reach in the hippocampus might be reduced resulting in each PV+ interneuron inhibiting a different number of postsynaptic cells in the local network, introducing variability in the synchronization of cell assemblies. Furthermore, recent work proposed that neuronal entrainment at γ-frequencies during development can impact neuronal morphology of cortical cells (Bitzenhofer et al., 2019). Thus, it is of great interest to explore the relationship between the fidelity of network activity at γ-frequency and neuronal morphology.

Surprisingly, given the established role of mitochondria in presynaptic release (Devine and Kittler, 2018; Sun et al., 2013), neither spontaneous inhibitory postsynaptic transmission nor PV+ interneuron mediated evoked inhibition were altered by the reduction of mitochondria in axonal boutons in Miro1^(Δ/Δ)^ (**Supplementary Fig 2**). Furthermore, these observations demonstrated that the changes in mitochondrial distribution associated with the impairments in Miro1-directed trafficking are not sufficient to alter the perisomatic inhibition applied by PV+ interneurons. The ability of PV+ interneurons to sustain inhibition and recover remained intact in the absence of Miro1 (**Supplementary Fig 3**). It is possible that the increased cellular concentration of mitochondria in PV+ interneurons ensures sufficient metabolic capacity to support their energetic demands. Indeed, the diffusion of ATP from mitochondria rich regions to boutons devoid of mitochondria seems to be sufficient to sustain the energetic requirements of synapses (Pathak et al., 2015). Further investigation is required to understand how the changes in Miro1-dependent mitochondrial positioning and enhanced axonal branching could be implicated in varying the strength and pattern of inhibition in the hippocampus.

Changes in parvalbumin interneuron excitability and network activity have been associated with alterations in behavior and emergence of neurological and neuropsychiatric disorders (Inan et al., 2016; Marín, 2012; Pelkey et al., 2017; Zou et al., 2016). Thus, we performed experiments to test exploratory locomotion, memory and anxiety in control and Miro1^(Δ/Δ)^ mice and assess whether the alterations in mitochondrial dynamics and axonal morphology were physiologically relevant by having an impact on behavior. Miro1^(Δ/Δ)^ animals presented no deficits in husbandry behavior, motor coordination and learning, short-term memory and spatial exploration when compared to their littermate controls (**Supplementary Fig 4**). Interestingly, Miro1^(Δ/Δ)^ animals exhibited anxiolytic behavior, demonstrated as increased time spent in the open arms of the elevated plus maze (EPM). This observation raises the interesting question about the role of Miro1-dependent mitochondrial dynamics and PV+ interneuron signaling in distinct brain areas involved in emotional behaviors such as the ventral hippocampus, amygdala, and prefrontal cortex (Janak and Tye, 2015). In conclusion, our findings demonstrate that the Miro1-directed spatiotemporal positioning of mitochondria in PV+ interneurons can modulate axon morphology, the frequency of hippocampal network oscillations at the γ-band range and potentially influence stress and emotional behaviors.

## ACKNOWLEDGMENTS

This work was supported by a PhD studentship from the Medical Research Council (MRC) to G.K (1405150) and grants from the MRC (MR/M024083/1) to P.C.S, the European Research Council grant 282430 (Fuelling Synapses) and the Lister Institute of Preventive Medicine to J.T.K.

## AUTHOR CONTRIBUTIONS

G.K and J.T.K conceptualized the project, designed experiments and wrote the paper. G.K performed live and fixed imaging, extracellular recordings, animal behavior experiments and analyzed all data. P.A and E.O.M designed the electrophysiological experiments and kindly provided recording equipment. P.A performed all intracellular recording experiments. M.P designed animal behavior experiments that were conducted in the facility provided by P.C.S. G.K and M.P performed the animal behavior experiments. N.F.H and G.L-D assisted in training and supervision. G.L-D designed animal breeding strategy. P.A, I.L.A-C and E.O.M wrote analysis scripts. G.K, P.A, M.P, N.F.H, B.R.S, G.L-D, P.C.S, E.O.M and J.T.K edited the paper.

---

**Supplementary Figure 1.**
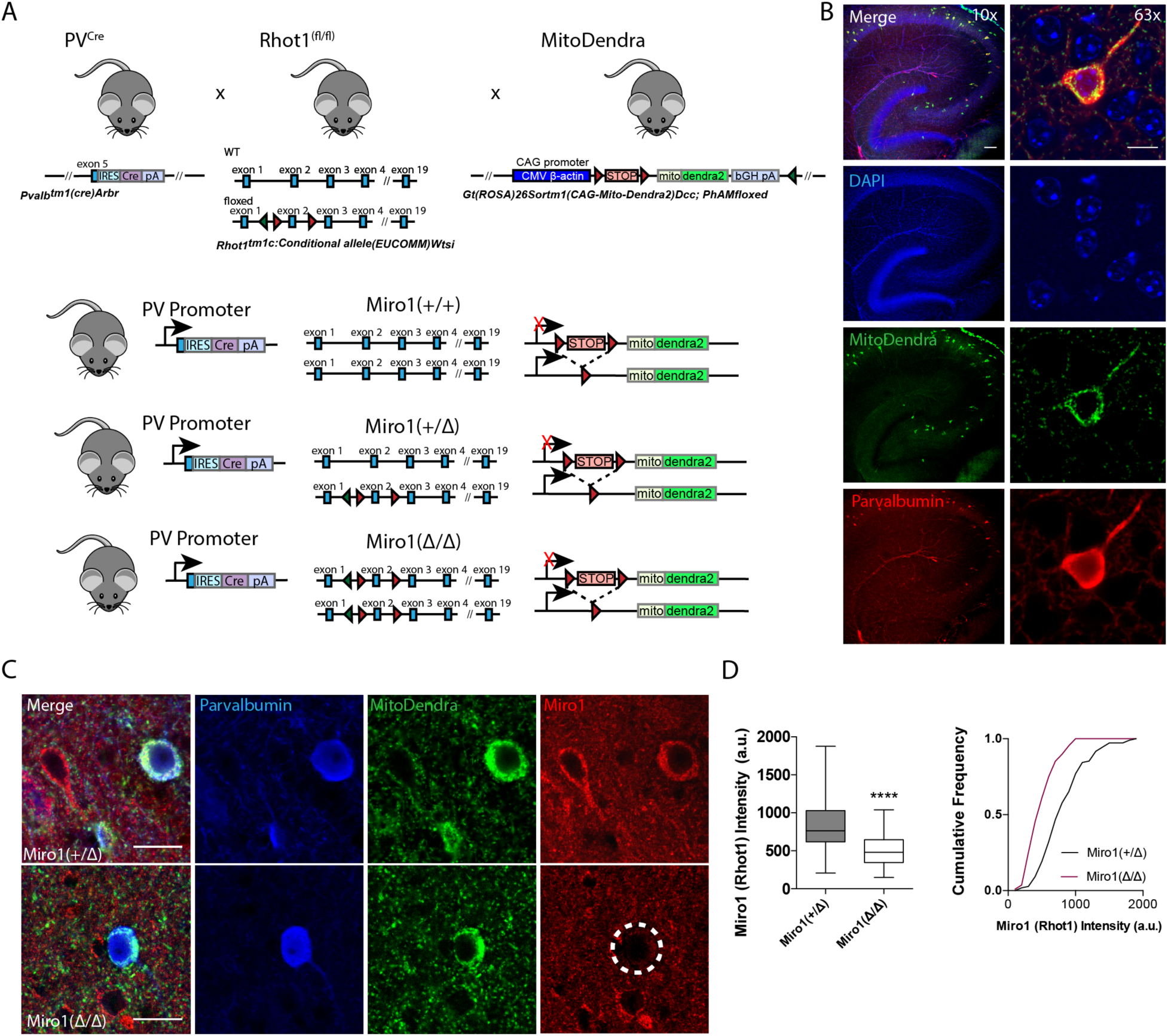
Generation of the PV^Cre^ *Rhot1*^(fl/fl)^ MitoDendra transgenic mouse line and conditional removal of Miro1 from MitoDendra-expressing parvalbumin interneurons. **A.** Schematic diagram of the genetic recombination for the expression of MitoDendra and simultaneous conditional removal of Miro1. When the Cre recombinase is expressed, under the PV+ promoter, the stop-floxed codon in the Rosa26 locus is excised allowing for the downstream expression of MitoDendra. Additionally, the second exon of the *Rhot1* gene is found between two loxP sites and therefore removed selectively in PV+ interneurons **B.** Cell-type specific expression of MitoDendra in PV+ interneurons. Left: Representative low magnification (10x) confocal image demonstrating the PV+ interneuron distribution in the acute brain slices of the hippocampus in the PV^Cre^ MitoDendra mouse. Scale Bar = 100 μm. Right: Representative high magnification (63x) image of a MitoDendra-expressing cell that is also PV immuno-positive. Scale Bar = 10 μm **C.** Loss of Miro1 from PV+ expressing interneurons. Representative confocal image from hemi-floxed control (Miro1^(+/Δ)^) and conditional knock-out (Miro1^(Δ/Δ)^) animals. Scale Bar = 10 μm. The fluorescent signal for Miro1 is specifically reduced in PV immuno-positive cells (white dotted circle shows the PV immuno-positive cell with low Miro1 fluorescence). **D.** Boxplot and cumulative frequency distribution of the *Rhot1* (Miro1) fluorescent intensity signal (n_Miro(+/Δ)_ = 109 neurons, 4 slices, 2 animals and n_Miro(Δ/Δ)_ = 109 neurons, 4 slices, 2 animals).

**Supplementary Figure 2.**
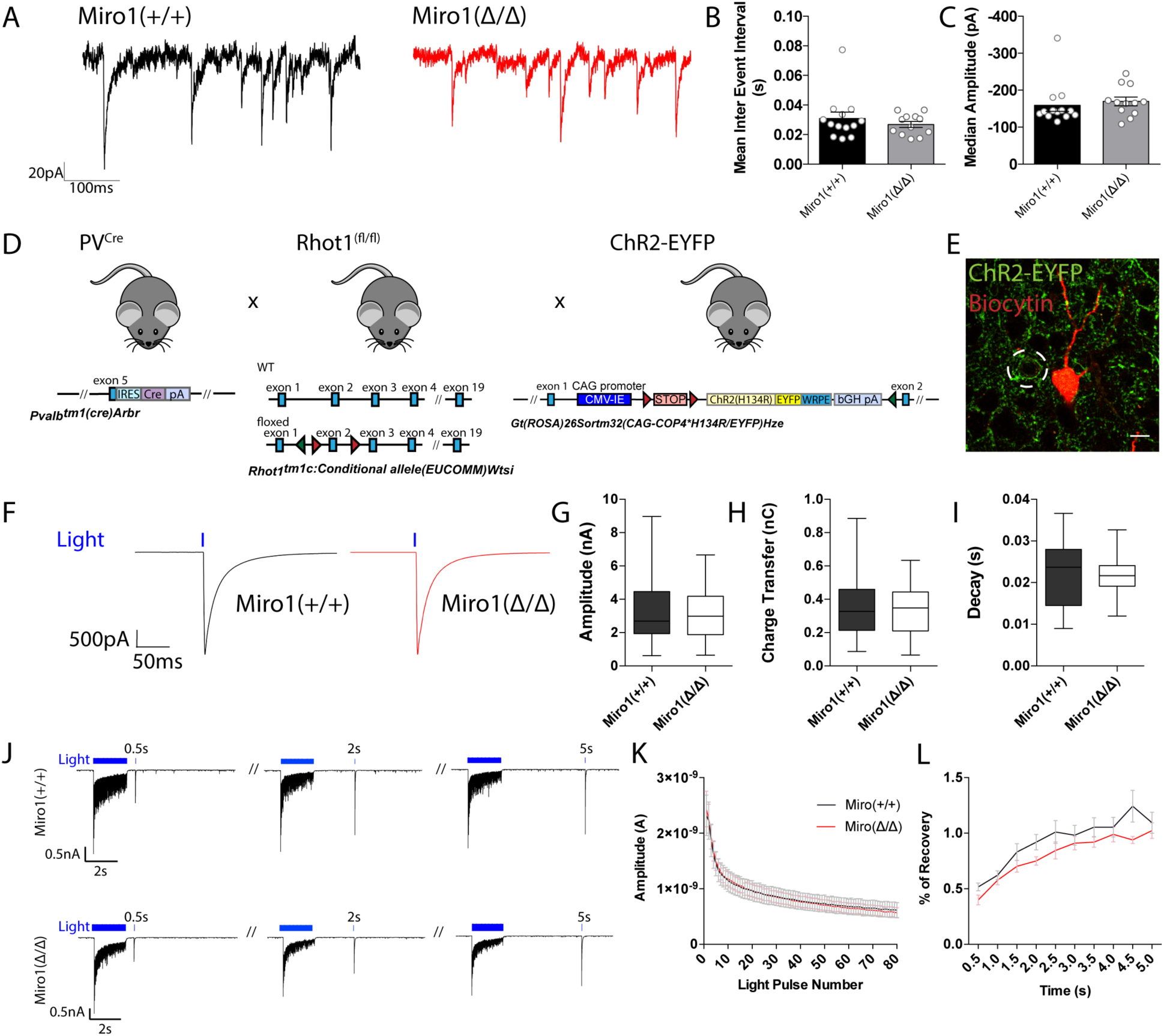
Miro1 knock-out does not alter spontaneous and evoked inhibitory synaptic transmission in the hippocampus. **A.** Representative electrophysiological traces of spontaneous inhibitory post-synaptic currents from Miro1^(+/+)^ (black) and Miro1^(Δ/Δ)^ (red) cells in the hippocampus. **B.** Quantification for the mean inter-event interval (IEI) **C.** Quantification for the median sIPSC amplitude. (n_Miro(+/+)_ = 13 recordings, 2 animals and n_Miro(Δ/Δ)_ = 12 recordings, 2 animals) **D.** Generation of the PV^Cre^ *Rhot1* ChR2-EYFP transgenic mouse line. Schematic diagram of the expression of ChR2-EYFP and simultaneous conditional removal of *Rhot1*. When the Cre recombinase is expressed, under the PV+ promoter, the stop-floxed codon is excised from the *Rosa26* locus allowing the downstream expression of ChR2-EYFP. Additionally, the second exon of the *Rhot1* gene is found between two loxP sites and also removed selectively in PV+ interneurons. **E.** Example of confocal image from a biocytin-filled recorded pyramidal cell in the hippocampus (red) in close proximity to a YFP+ PV+ interneuron (green). Scale bar = 10 μm. **F.** Representative traces from light evoked inhibitory postsynaptic current (eIPSC) in Miro1^(+/+)^ (black) and Miro1^(Δ/Δ)^ (red) cells in acute brain slices (n_Miro(+/+)_ = 23 recordings, 4 animals and n_Miro(Δ/Δ)_ = 23 recordings, 4 animals). **G.** Boxplot for the quantification of peak amplitude. **H.** Boxplot for the quantification of charge transfer. **I.** Boxplot for the quantification of decay. **J.** Control and conditional knock-out cells can sustain inhibition and recover with similar rates after long-lasting photostimulation. Example traces from the inhibitory responses pyramidal cells received in Miro1^(+/+)^ and Miro1^(Δ/Δ)^ slices during light train stimulation (40 Hz for 2s; 1 ms pulse width) **K.** Mean amplitude of each peak during the light train stimulation (n_Miro(+/+)_ = 21 recordings, 4 animals and n_Miro(Δ/Δ)_ = 24 recordings, 4 animals). **L.** Quantification of the percentage recovery after light stimulation of all cells at increasing time intervals from the end of the light train (n_Miro(+/+)_ = 21 recordings, 4 animals and n_Miro(Δ/Δ)_ = 18 recordings, 3 animals).

**Supplementary Figure 3.**
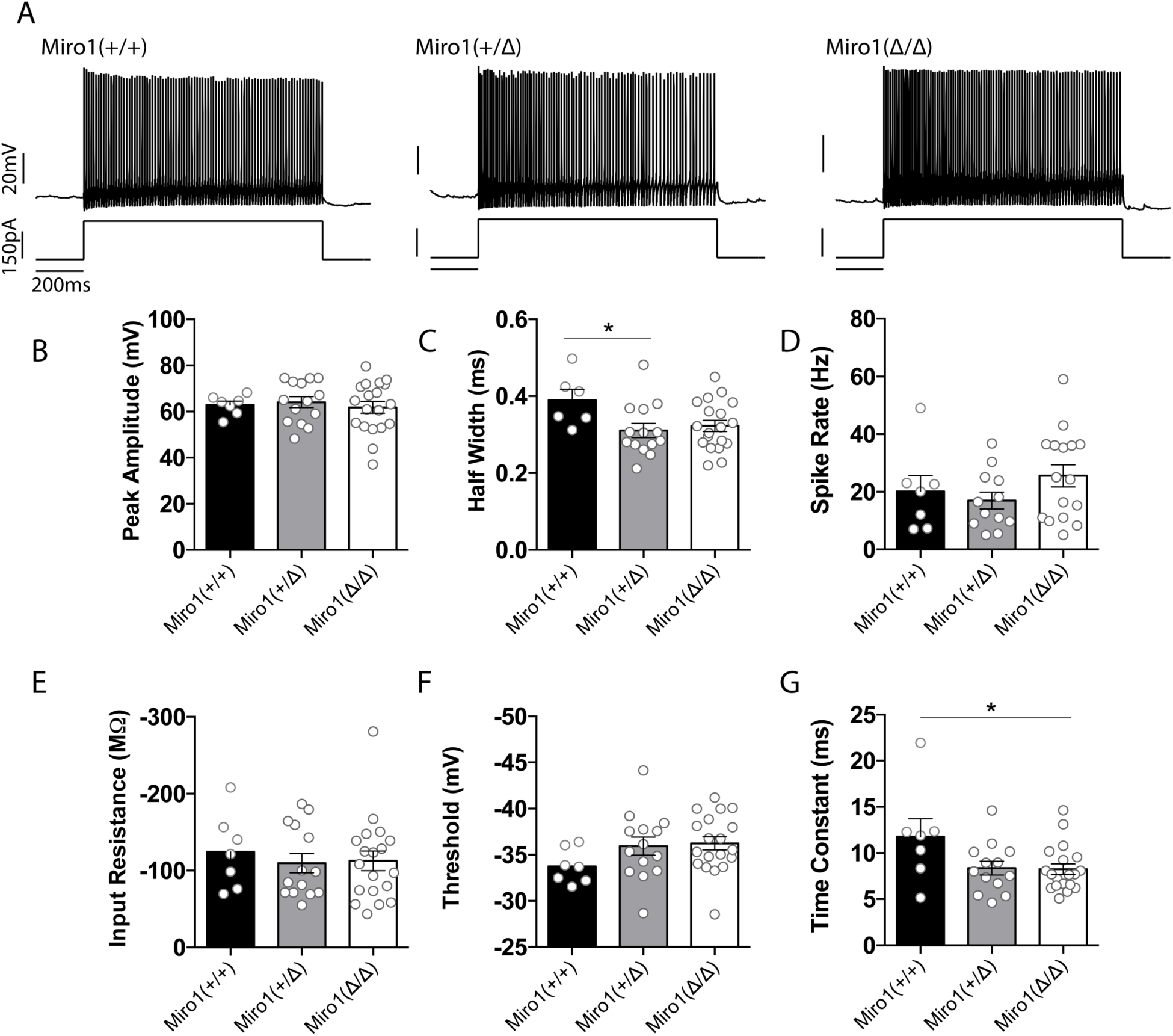
Intrinsic properties of PV+ interneurons in the absence of Miro1. **A.** Representative traces of the firing patterns of Miro1^(+/+)^, Miro1^(+/Δ)^ and Miro1^(Δ/Δ)^ cells in response to 200 pA current injection (n_Miro(+/+)_ = 2 animals, n_Miro(+/Δ)_ = 4 animals, n_Miro(Δ/Δ)_ = 5 animals). **B.** Quantification of the action potential peak amplitude **C.** Quantification of the action potential half-width. **D.** Quantification of Spike Rate at RheoBase +40 pA. Cells that fire less than 5 spikes were excluded from the quantification. **E.** Quantification of input resistance **F.** Quantification of threshold to fire **G.** Quantification of time constant.

**Supplementary Figure 4.**
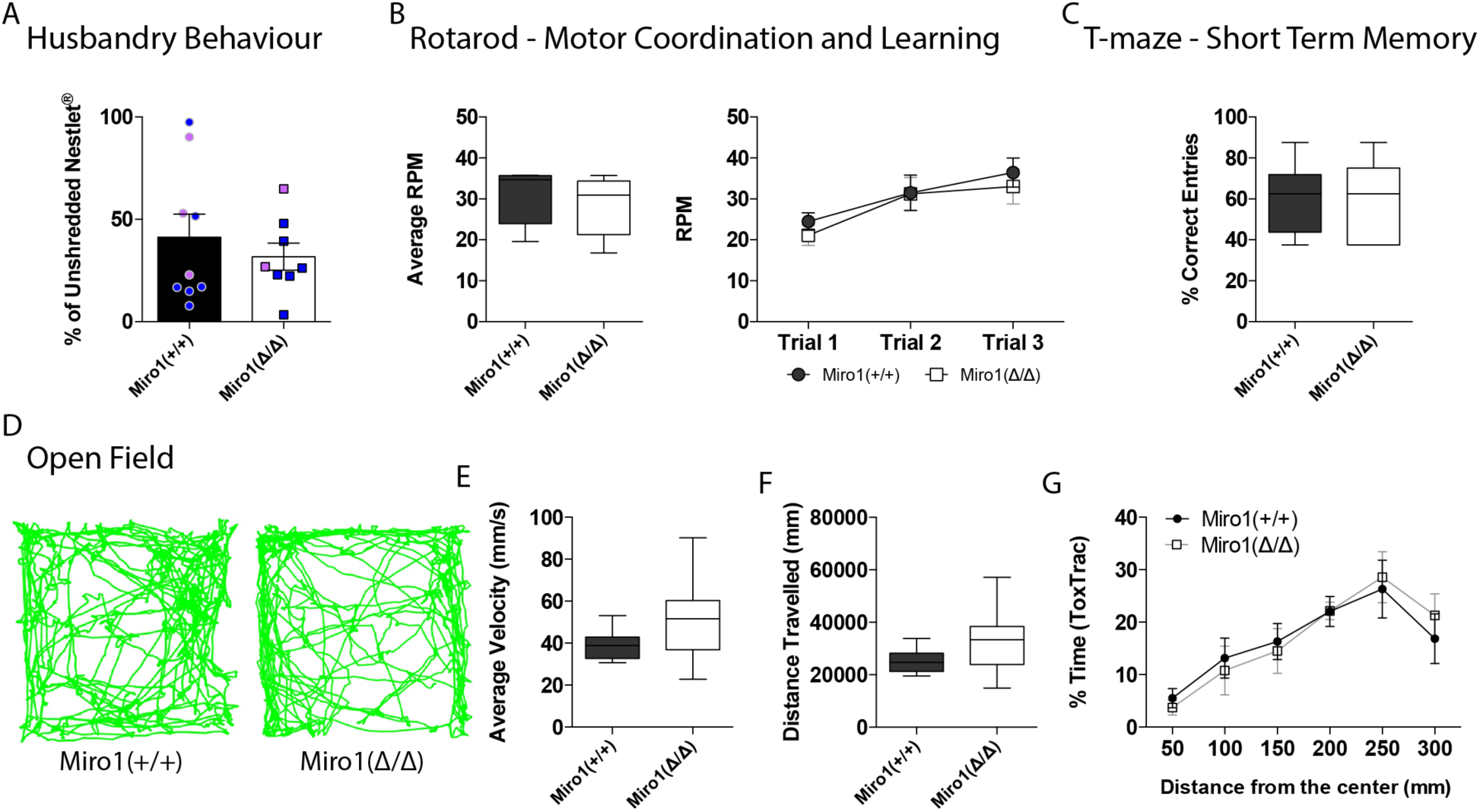
Loss of Miro1 does not affect husbandry behavior, motor coordination, short term memory and spatial exploration. **A.** Assessment of husbandry behavior based on the amount of shredded and unshredded Nestlet between Miro1^(+/+)^ and Miro1^(Δ/Δ)^ animals. Bar chart shows the quantification of the percentage of unshredded Nestlet. Blue points represent values from male mice and pink points represent values from female mice (n_Miro(+/+)_ = 9 animals, n_Miro(Δ/Δ)_ = 8 animals). **B.** Assessment of motor coordination using the rotarod. The animal is placed on the revolving rot until it falls and the speed at which it falls is registered to calculate the revolutions per minute (RPM) indicated by the box plot (n_Miro(+/+)_ = 5 animals, n_Miro(Δ/Δ)_ = 5 animals). **C.** Assessment of short-term memory using the spontaneous alternation T-maze behavioral paradigm. The animal starts every trial at position marked as start, the animal is allowed to make a decision at the end of the maze and is kept in that arm for 30 seconds. A correct entry is considered when the animal makes the opposite decision from the previous trial. A wrong entry is when the animals makes the same decision as in the previous trial. Box plot shows the quantification of the percentage of correct entries. (n_Miro(+/+)_ = 8 animals, n_Miro(Δ/Δ)_ = 7 animals). **D.** Assessment of general exploration in an open field. Example trajectories (green) of Miro1^(+/+)^ and Miro1^(Δ/Δ)^ animals. **E.** Box plot for the quantification of the average velocity. **F.** Box plot for the quantification of the distance travelled. **G.** Quantification of the percentage time spent at distances away from the centre of the box (n_Miro(+/+)_ = 9 animals, n_Miro(Δ/Δ)_ = 9 animals).

## EXPERIMENTAL MODEL

### Animals

All experimental procedures were carried out in accordance with institutional animal welfare guidelines and licensed by the UK Home Office in accordance with the Animals (Scientific Procedures) Act 1986. Animals were maintained under controlled conditions (temperature 20 ± 2°C; 12-hour light-dark cycle). Animals of either sex were used for all experiments. The PV^Cre^ line (Stock Number 008069) has been previously described in (Hippenmeyer et al., 2005). The *Rhot1* transgenic line (Rhot1tm1a (EUCOMM)Wtsi) was obtained from the Wellcome Trust Sanger Institute (MBTN EPD0066 2 F01) and the floxed mouse has been previously generated using the Flp recombination strategy described here (López-Doménech et al., 2016). Briefly, the exon 2 of *Rhot1* gene (chromosome 11) is flanked by two LoxP sites and Cre recombination results in the deletion of exon 2. The stop-floxed MitoDendra line (*B*6; 129*S Gt*(*ROSA*)26*Sortm*1(*CAGCOX*8*A*/*Dendra*2)*Dcc*/*J*) (Stock number 018385) and the stop-floxed ChR2-EYFP (*B*6;129*SGt*(*ROSA*)26*Sortm*32(*CAGCOP*4\**H*134*R*/*EYFP*)*Hze*/*J*) (Stock number 012569) were also obtained from The Jackson Laboratory. The MitoDendra line has been previously described in (Pham et al., 2012) and the ChR2-EYFP line in (Madisen et al., 2012). Experimental animals were generated as a result of the following crosses: PV^Cre+/-^ *Rhot1*^fl/fl^ x PV^Cre-/-^ *Rhot1*^fl/fl^, PV^Cre+/+^ *Rhot1*^+/+^ MitoDendra^+/+^ x PV^Cre+/+^ *Rhot1*^+/+^ MitoDendra^+/+^, PV^Cre+/+^ *Rhot1*^+/fl^ MitoDendra^+/+^ x PV^Cre+/+^ *Rhot1*^fl/fl^ MitoDendra^+/+^ and PV^Cre+/+^ *Rhot1*^+/fl^ ChR2-EYFP^+/+^ x PV^Cre+/+^ *Rhot1*^+/fl^ ChR2-EYFP^+/+^. Genotyping was carried out following Sanger recommended procedures on ear biopsies for adult mice and tail biopsies for neonatal mice (<P10).

## METHOD DETAILS

### Hippocampal Brain Slice Preparations

#### Acute brain slices

Adult mice (>P60) were anesthetized using 4% isoflurane followed by decapitation, and the brains were extracted in warm (30-35°C) sucrose solution [40 mM NaCl, 3 mM KCL, 7.4 mM MgSO_4_.7H_2_O, 150 mM sucrose, 1 mM CaCl_2_, 1.25 mM NaH_2_PO_4_, 25 mM NaHCO_3_ and 15 mM glucose; Osmolality 300 ± 10 mOsmol/Kg]. Horizontal hippocampal slices (350 μm thick) were cut using a vibratome (Leica VT1200S) and were placed in an interface chamber containing warm artificial cerebrospinal fluid (aCSF) [126 mM NaCl, 3.5 mM KCl, 2 mM MgSO_4_.7H_2_O, 1.25 mM NaH_2_PO_4_, 24 mM NaHCO_3_, 2 mM CaCl_2_ and 10 mM glucose; Osmolality 300 ± 10 mOsmol/Kg]. All solutions were bubbled with carbogen gas [95% O_2_/ 5% CO_2_].

#### Organotypic brain slices

*Neonatal* mice (P6-8) were sacrificed by cervical dislocation followed by decapitation. The brains were extracted in ice-cold dissection medium [487.5 ml Earle’s Balanced Salt Solution (EBSS) and 12.5 ml of 25 mM HEPES]. 300 μm thick transverse hippocampal slices were cut using a vibratome (Leica VT1200S) in ice-cold dissection medium. Organotypic slices were prepared using the Stoppini interface method as described in (De Simoni and Yu, 2006; Stephen et al., 2015; Stoppini et al., 1991). Briefly, the slices were kept on sterile 0.45 μm Omnipore membrane inserts (Millipore, cat no. FHLC01300) in an incubator (37°C, 95% O_2_/ 5% CO_2_) for at least 6 days in culture media [47% MEM + GlutaMAX, 25% horse serum, 25% EBSS supplemented with 20 mM HEPES, 1.44% of 45% glucose, 1.06% penicillin/streptomycin with 16% nystatin, and 0.5% 1 M Tris solution] prior to imaging. The media were changed on the day of slicing and half of the media were replaced with fresh media every 3-4 days.

### Microscopy

#### Fixed Confocal Imaging

Confocal images (1024 x 1024) were acquired on a Zeiss LSM700 upright confocal microscope using the 10x air, 20x water and 63x oil objective and digitally captured using the default LSM acquisition software. For analysis, 2-3 zoomed regions of the hippocampus were imaged with the 2x zoom. For the quantification, these regions were averaged and represented as one value. Acquisition settings and laser power were kept constant within experiments. For neuronal reconstructions, ∼150 μm thick confocal stacks were captured using the 20x objective, with z-steps of 1 μm.

#### Live Two-photon Imaging

Organotypic slices were live-imaged using the 20x and 60x water objectives on a Zeiss LSM700 upright two photon microscope equipped with a MaiTai Ti:Saphire Laser (Spectra-Physics). The slices were transferred to a recording chamber, perfused with aCSF [2 mM CaCl_2_, 2.5 mM KCl, 1 mM MgCl_2_, 10 mM D-glucose, 126 mM NaCl, 24 mM NaHCO_3_, 1 mM NaH_2_PO_4_] bubbled with carbogen gas and heated between 35-38°C at a constant perfusion (∼2 ml/min). The excitation wavelength was set at 900nm and the rate of image acquisition was 1 frame / 5 seconds for 500 seconds (100 frames per movie).

### Tissue Processing and Labeling

#### Brain Harvesting

Adult animals (>P60) were sacrificed by cervical dislocation or CO_2_ exposure. The brains were dropped-fixed in 4% paraformaldehyde (PFA) in sucrose solution overnight at 4°C. The brains were then cryo-protected in 30% sucrose/1X Phosphate Buffered Saline (PBS) [1.37 mM NaCl, 2.7 mM KCl, 10 mM Na_2_HPO_4_, 2 mM KH_2_PO_4_] solution overnight at 4°C before freezing at −80°C. Hemi-floxed and conditional knock-out animals expressing the MitoDendra fluorophore were anaesthetized with isoflurane and transcardially perfused with ice-cold 4% PFA to maintain mitochondrial morphology. The frozen brains were embedded in tissue freezing compound (OCT) and 30 μm coronal brain slices were serially cryosected using a Cryostat (Bright Instruments). After live-imaging and electrophysiological recordings, brain slices were fixed in 4% PFA/sucrose solution overnight at 4°C. Slices were either kept in PBS at 4°C for short-term storage or at −20°C in cryoprotectant solution [30% glycerol, 30% ethylene glycol, 40% 1X PBS] for long-term storage.

#### Immunohistochemistry

Free floating sections were washed with 1X PBS and permeabilized for 4-5 hours in block solution [1X PBS, 10% horse serum supplemented with 0.02% sodium azide, 3% (w/v) Bovine Serum Albumin (BSA), 0.5% Triton X-100 and 0.2 M glycine]. The slices were further blocked overnight with an added purified goat anti-mouse Fab-fragment (50 μg/ml, Jackson Immunoresearch) for reducing endogenous background. The sections were then incubated with primary antibody diluted in block solution overnight at 4°C. The following primary antibodies were used: parvalbumin (mouse, 1:500, Millipore MAB1572), COX-IV (rabbit, 1:500, Abcam ab16056) and Rhot1 (rabbit, 1:100, Atlas HPA010687). Slices were washed 4-5 times in 1X PBS over 2 hours and then incubated for 3-4 hours with secondary antibody in block solution (1:500-1:1000) at room temperature. The secondary antibodies used were the donkey anti-mouse Alexa Fluor 488 (Jackson ImmunoResearch 715-545-151), goat anti-rabbit Alexa Fluor 555 (Thermo Fisher Scientific A-21430) and donkey anti-rabbit Alexa Fluor 568 (Thermo Fisher Scientific A-10042). The slices were then washed 4-5 times in PBS for 2 hours and mounted onto glass slides using Mowiol mounting media.

#### Biocytin Labeling

Biocytin-filled slices were fixed in 4% PFA solution after intracellular recordings and kept overnight at 4°C. The slices were washed with 1X PBS 3-4 times and permeabilized with 0.3%-Triton 1X PBS for 4-5 hours. Streptavidin conjugated to Alexa Fluor 555 (Invitrogen S32355) in PBS-T 0.3% (1:500) was added and the slices were kept overnight at 4°C. The slices were then washed 4-5 times in 1X PBS over 2 hours. The slices were placed on glass slides and a coverslip was mounted on top using Dako Fluorescent mounting medium.

### Image Analysis

#### Mitochondrial Trafficking

The image sequences were subjected to alignment (stackreg) if necessary, background subtraction (rolling ball radius = 50 pixels) and filtering (smooth filter). Moving mitochondria were visually identified. Mitochondria were manually tracked between each frame using MTrackJ (Meijering et al., 2012) on Fiji, which provided track statistics. Mitochondrial movement was usually accompanied by brief periods of immobility so data were omitted from the velocity (μm.s^−1^) calculations when a mitochondrion was immobile for a period longer than 10 seconds. Mitochondria were considered mobile if the distance covered was longer than 2 μm in 5 minutes.

#### Fluorescent Intensity

The fluorescence intensities of COX-IV and *Rhot1* were quantified on Fiji. The signal from the parvalbumin channel was thresholded using the default settings, the parvalbumin cell body was selected using the wand tool and a mask was generated that was then superimposed on the channels of interest to selectively record the fluorescent signal within the masked region.

#### Neuronal Reconstruction

The biocytin signal in acute brain slices was used to manually reconstruct parvalbumin interneurons in Neuromantic (Myatt et al., 2012) and generate .SWC files for further analysis.

#### Sholl Analysis

3D Sholl analysis was performed on the .SWC file using the Matlab script described in (Madry et al. 2018). To perform the 3D-MitoSholl, the volume of the .SWC file was filled out and a stack-mask was generated in the Simple Neurite Tracer Plugin on Fiji (Longair et al., 2011). To isolate mitochondria selectively in parvalbumin interneurons, the background was subtracted (rolling ball radius = 50 pixel) and the median filter was applied prior to binarization. The stack of the mitochondrial distribution in individual cells was generated by adding the stack-mask to the mitochondrial channel using the “AND” function of the Image Calculator in Fiji. 3D-MitoSholl analysis was performed using a custom Matlab script which quantified the number of MitoDendra pixels within each sholl ring, radiating out from the soma at 1 μm intervals.

#### Proximity Analysis

High magnification confocal stacks of the parvalbumin interneuron axon were acquired from 350 μm slices for calculating the minimum distance between mitochondria and boutons/branch points. The minimum distance between the boutons and the mitochondria was measured on max-projected images. The biocytin signal was used as a mask to isolate mitochondria in the axon only. The mitochondrial image was binarized using the default method. The (x,y) coordinates of the boutons were manually specified and the minimum distance between the bouton coordinates and the closest mitochondrion was calculated using a custom Matlab script, which calculated the Euclidean distance between the x and y coordinates and the first encountered binary pixel. The minimum distance between branch points and mitochondria was performed on 3D confocal stacks. The mitochondrial confocal stack was binarised using the default method). The (x,y,z) coordinates of the branch points were specified and the minimum distance between the branch points coordinates and the closest mitochondrion was calculated using a custom Matlab script, which calculated the Euclidean distance between the x,y,z coordinates and the first pixel encountered.

### Electrophysiology

#### Extracellular Recordings

Local field potentials (LFPs) were recorded in an interface recording chamber as described in (Antonoudiou et al., 2020). Briefly, an extracellular borosilicate glass electrode (tip resistance 1-5 MΩ) was filled with aCSF [126 mM NaCl, 3.5 mM KCl, 2 mM MgSO_4_.7H_2_O, 1.25 mM NaH_2_PO4, 24 mM NaHCO_3_, 2 mM CaCl_2_ and 10 mM glucose; Osmolality 300 ± 10 mOsmol/Kg] and was placed in hippocampal area CA3. γ-oscillations (20-80Hz) were induced by perfusion of carbachol (5 μM) in carbogen-bubbled aCSF. Data were acquired using 10 kHz sampling rate and amplified x10 by Axoclamp 2A (Molecular Devices). The signal was further amplified x100 and low-pass filtered at 1kHz (LPBF-48DG, NPI Electronic). The signal was digitized at 5 kHz by a data acquisition board (ITC-16, InstruTECH) and recorded on Igor Pro 6.3 software (Wavemetrics).

#### Intracellular Recordings

Intracellular recordings were performed in a submerged chamber (32-33°C) using borosilicate glass pipettes (5-12 MΩ). Data were acquired through the MultiClamp 700B amplifier (Molecular Devices) and digitized at 10 kHz (ITC-18, InstruTECH). Acquisition of electrophysiological signals was performed using Igor Pro 6.3 (Wavemetrics). The signals were low-pass filtered at 10 kHz and 3 kHz for current-clamp and voltage-clamp, respectively. For optogenetic experiments, filtered white LED (460 ± 30 nm, THOR labs) was delivered via epi-illumination through a 60x objective and was used to activate ChR2. For whole cell current clamp recordings, pipettes were filled with internal solution [110 mM K-Gluconate, 40 mM HEPES, 2 mM ATP-Mg, 0.3 mM GTP-NaCl, 4 mM NaCl, 3-4 mg/ml biocytin (Sigma); pH∼7.2; Osmolality 270-290 mOsmol/Kg]. After break through, the bridge balance was adjusted to compensate for electrode access. Hyperpolarizing and depolarizing square current pulses were applied in order to quantify intrinsic properties of the recorded neuron. sEPSCs on parvalbumin interneurons were recorded in voltage clamp mode at −70 mV, using the current clamp internal solution. IPSCs on pyramidal cells were recorded in voltage clamp mode with pipettes filled with internal solution [137 mM CsCl, 5 mM NaCl, 10 mM HEPES, 0.1 mM EGTA, 2 mM ATP-Mg, 0.3 mM GTP-Na, 3-4 mg/ml biocytin]. These experiments were done on two batches of animals. For evoked inhibitory post-synaptic currents (eIPSCs) the cells were held at −40 mV. This was to prevent spiking during light illumination in the absence of QX-314. In the second set of recordings, 5 mM QX-314 was added and the voltage was still held at −40 mV. The recordings between the first and second batches were comparable and therefore pooled together and quantified as one dataset. For optogenetic experiments, aCSF was supplemented with 3 mM kynurenic acid. After breakthrough the cell and electrode capacitance were compensated in Multiclamp. Series resistance (RS) compensation was performed to 65%. For optogenetic experiments, a power plot using increasing light intensities was generated to decide the level of LED voltage to be used. The LED voltage that elicited 90% of the maximal response was used for stimulations. The LED output range used for power plot was between 0-1.53 mW.

#### Electrophysiological Analysis

In order to characterize and analyze the γ-oscillations, we calculated power spectra as the normalized magnitude square of the FFT (Igor Pro 6.3). The 50 Hz frequency was not included in the analysis to exclude the mains noise. The oscillation amplitude was quantified by measuring the peak of the power spectrum (peak power) and the area below the power spectrum plot (power area) within the γ-band range (20-80 Hz). The peak frequency of the oscillation was the frequency at which the peak of the power spectrum occurred in the γ-band range. For testing rhythmicity, autocorrelation was computed in Igor Pro 6.3. The peak power, power area and peak frequency were calculated for a period of 300 seconds and compared between control, hemi-floxed and cKO mice. The power spectra were also calculated over a period of 300 seconds and a gaussian curve fit was performed using Igor Pro 6.3. This was done in order to assess the width of the power spectrum distribution. For spontaneous post-synaptic current (sPSC) detection, custom written procedures were used in Igor Pro 6.3. The traces were first low-pass filtered at 1 kHz. sPSCs were first detected using an initial threshold of 3 pA. The detected events that were smaller than 5x standard deviation of noise were excluded from further analysis. The traces were then visually inspected for correct peak identification. The traces were averaged, and the median inter-event interval and peak amplitude were obtained for each cell. For the recovery experiments, the percentage of recovery was normalized to the amplitude of the first peak in the train, for every train and was presented as an average of the 10 trains.

### Animal Behavior Testing

#### Nesting Assesment

The assessment husbandry behavior using Nestlets was performed as described in (Deacon, 2006). Briefly, one hour before the dark cycle, mice were transferred in single cages containing wood-chip bedding but no other environmental enrichment. One Nestlet (5 x 5 cm) was placed in each cage. The following morning the shredded and unshredded Nestlets were collected and weighted.

#### Open field

The open field apparatus consisted of a 45 x 45 x 45 cm box open at the top. Mice were placed in the center of the arena and were recorded for 10 minutes via a ceiling-fixed video camera. The movies were analyzed using ToxTrac, an automated open-source executable software (Rodriguez et al., 2017). Detection settings were set to 10 for threshold, 100 for minimum object size and 1000 for maximum object size and the default tracking settings were used.

#### Rotarod

The assessment of motor coordination and learning was performed similar to (Deacon, 2013). Briefly, mice were placed on a revolving rotarod treadmill (Med Associates) with a starting speed of 4 revolutions per minute (RPM) and an acceleration rate of about 7 RPM/min. Each mouse was subjected to a 5 minute trial, repeated 2 more times after a 10 minute gap. To calculate the RPM, we recorded the speed at which the mouse fell from the revolving rod using this formula: RPM = [(End Speed - Start Speed) / 300] x (Seconds Run) + Start Speed.

#### T-Maze

The assessment of spontaneous alternation on the T-maze was performed according to (Deacon and Rawlins, 2006). The T-maze apparatus consisted of three arms with 27 x 7 x 10cm dimensions. A set of three sliding guillotine doors was used to separate the entrance of each arm. At the beginning of the experiment all of the doors were raised, except for the one located in front of the starting point. Each mouse was allowed 8 consecutive trials. Each trial consisted of individually placing each animal at the entrance of the main arm and allowing it to freely run and chose an arm. After the mouse entered an arm, the guillotine door was pushed down and the animal was confined in the chosen arm for 30 seconds. The mouse was then placed in the starting point for 30 seconds before the beginning of the next trial. Each trial lasted less than 90 seconds. When a mouse did not choose an arm within 90 seconds it was considered as a NO-GO and the trial was restarted. Animals were excluded from the quantification when there were more than 3 NO-GOs.

#### Elevated Plus Maze

The elevated plus maze was placed 40 cm above the ground and consisted of four 30 x 5 cm arms, two of which were surrounded by additional 15 cm high walls. Mice were placed on the boundary between the open arm and the center, facing the center. Mice were recorded for 5 minutes by a ceiling-fixed video camera and the movies were analyzed using ToxTrac (Rodriguez et al., 2017).

### Statistical Analysis and Blinding

Statistical analysis was performed in Prism 6, Excel and Igor Pro. Datasets were tested for fitting in a Gaussian distribution using the D’Agostino-Pearson omnibus and the Shapiro-Wilk normality test. When the distribution was normal, unpaired t-test was performed. The F-test was used to compare variances. When the variance was significantly different, the unpaired t-test with Welch’s correction was performed. For cumulative distributions, the Kolmogorov-Smirnov (KS) test and multiple t-test per row were performed. For non-normally distributed data, the Mann-Whitney (MW) test was applied. For comparing multiple groups, ordinary one-way ANOVA and Tukey’s multiple comparisons test with a single pooled variance were used. Statistical outliers were identified using the ROUT method and removed (Motulsky and Brown, 2006). Significance of p < 0.05 is represented as *, p < 0.01 as **, p < 0.001 as *** and p < 0.0001 as ****. The errors in all bar charts represent the standard error of the mean (sem). Brain tissue processing, staining, acquisition and analysis were performed blinded. Live imaging acquisition and analysis were performed blinded for littermate hemi-floxed and conditional knock-out animals. Electrophysiological recording analysis was performed blinded.

## REFERENCES

Adams, D.L., Economides, J.R., and Horton, J.C. (2015). Co-localization of glutamic acid decarboxylase and vesicular GABA transporter in cytochrome oxidase patches of macaque striate cortex. Vis Neurosci 32, E026.

Akam, T., and Kullmann, D.M. (2010). Oscillations and filtering networks support flexible routing of information. Neuron 67, 308–320.

Antonoudiou, P., Tan, Y.L., Kontou, G., Upton, A.L., and Mann, E.O. (2020). Parvalbumin and somatostatin interneurons contribute to the generation of hippocampal gamma oscillations. The Journal of Neuroscience. doi: 10.1523/JNEUROSCI.0261-20.2020.

Attwell, D., and Laughlin, S.B. (2001). An energy budget for signaling in the grey matter of the brain. J. Cereb. Blood Flow Metab. 21, 1133–1145.

Barnes, S.A., Pinto-Duarte, A., Kappe, A., Zembrzycki, A., Metzler, A., Mukamel, E.A., Lucero, J., Wang, X., Sejnowski, T.J., Markou, A., et al. (2015). Disruption of mGluR5 in parvalbumin-positive interneurons induces core features of neurodevelopmental disorders. Mol. Psychiatry 20, 1161–1172.

Bartos, M., Vida, I., and Jonas, P. (2007). Synaptic mechanisms of synchronized gamma oscillations in inhibitory interneuron networks. Nat. Rev. Neurosci. 8, 45–56.

Birsa, N., Norkett, R., Higgs, N., Lopez-Domenech, G., and Kittler, J.T. (2013). Mitochondrial trafficking in neurons and the role of the Miro family of GTPase proteins. Biochem. Soc. Trans. 41, 1525–1531.

Bitzenhofer, S.H., Poepplau, J.A., Chini, M., Marquardt, A., and Hanganu-Opatz, I. (2019). Activity-dependent maturation of prefrontal gamma oscillations sculpts cognitive performance. BioRxiv.

Buzsáki, G., and Wang, X.-J. (2012). Mechanisms of gamma oscillations. Annu. Rev. Neurosci. 35, 203–225.

Cardin, J.A., Carlén, M., Meletis, K., Knoblich, U., Zhang, F., Deisseroth, K., Tsai, L.-H., and Moore, C.I. (2009). Driving fast-spiking cells induces gamma rhythm and controls sensory responses. Nature 459, 663–667.

Chada, S.R., and Hollenbeck, P.J. (2003). Mitochondrial movement and positioning in axons: the role of growth factor signaling. J. Exp. Biol. 206, 1985–1992.

Chada, S.R., and Hollenbeck, P.J. (2004). Nerve growth factor signaling regulates motility and docking of axonal mitochondria. Curr. Biol. 14, 1272–1276.

Chattopadhyaya, B., Di Cristo, G., Higashiyama, H., Knott, G.W., Kuhlman, S.J., Welker, E., and Huang, Z.J. (2004). Experience and activity-dependent maturation of perisomatic GABAergic innervation in primary visual cortex during a postnatal critical period. J. Neurosci. 24, 9598–9611.

Chattopadhyaya, B., Di Cristo, G., Wu, C.Z., Knott, G., Kuhlman, S., Fu, Y., Palmiter, R.D., and Huang, Z.J. (2007). GAD67-mediated GABA synthesis and signaling regulate inhibitory synaptic innervation in the visual cortex. Neuron 54, 889–903.

Courchet, J., Lewis, T.L., Lee, S., Courchet, V., Liou, D.-Y., Aizawa, S., and Polleux, F. (2013). Terminal axon branching is regulated by the LKB1-NUAK1 kinase pathway via presynaptic mitochondrial capture. Cell 153, 1510–1525.

De Simoni, A., and Yu, L.M.Y. (2006). Preparation of organotypic hippocampal slice cultures: interface method. Nat. Protoc. 1, 1439–1445.

Deacon, R.M.J. (2006). Assessing nest building in mice. Nat. Protoc. 1, 1117–1119.

Deacon, R.M.J. (2013). Measuring motor coordination in mice. J. Vis. Exp. e2609.

Deacon, R.M.J., and Rawlins, J.N.P. (2006). T-maze alternation in the rodent. Nat. Protoc. 1, 7–12.

Devine, M.J., and Kittler, J.T. (2018). Mitochondria at the neuronal presynapse in health and disease. Nat. Rev. Neurosci. 19, 63–80.

Freund, T.F., and Buzsáki, G. (1996). Interneurons of the hippocampus. Hippocampus.

Fries, P. (2015). Rhythms for Cognition: Communication through Coherence. Neuron 88, 220–235.

Galow, L.V., Schneider, J., Lewen, A., Ta, T.-T., Papageorgiou, I.E., and Kann, O. (2014). Energy substrates that fuel fast neuronal network oscillations. Front. Neurosci. 8, 398.

Glausier, J.R., Roberts, R.C., and Lewis, D.A. (2017). Ultrastructural analysis of parvalbumin synapses in human dorsolateral prefrontal cortex. J. Comp. Neurol. 525, 2075–2089.

Gulyás, A.I., Buzsáki, G., Freund, T.F., and Hirase, H. (2006). Populations of hippocampal inhibitory neurons express different levels of cytochrome c. Eur. J. Neurosci. 23, 2581–2594.

Guo, X., Macleod, G.T., Wellington, A., Hu, F., Panchumarthi, S., Schoenfield, M., Marin, L., Charlton, M.P., Atwood, H.L., and Zinsmaier, K.E. (2005). The GTPase dMiro is required for axonal transport of mitochondria to Drosophila synapses. Neuron 47, 379–393.

Hájos, N., Pálhalmi, J., Mann, E.O., Németh, B., Paulsen, O., and Freund, T.F. (2004). Spike timing of distinct types of GABAergic interneuron during hippocampal gamma oscillations in vitro. J. Neurosci. 24, 9127–9137.

Hippenmeyer, S., Vrieseling, E., Sigrist, M., Portmann, T., Laengle, C., Ladle, D.R., and Arber, S. (2005). A developmental switch in the response of DRG neurons to ETS transcription factor signaling. PLoS Biol. 3, e159.

Hirokawa, N., Niwa, S., and Tanaka, Y. (2010). Molecular motors in neurons: transport mechanisms and roles in brain function, development, and disease. Neuron 68, 610–638.

Howard, M.W., Rizzuto, D.S., Caplan, J.B., Madsen, J.R., Lisman, J., Aschenbrenner-Scheibe, R., Schulze-Bonhage, A., and Kahana, M.J. (2003). Gamma oscillations correlate with working memory load in humans. Cereb. Cortex 13, 1369–1374.

Hu, H., Gan, J., and Jonas, P. (2014). Interneurons. Fast-spiking, parvalbumin^+^ GABAergic interneurons: from cellular design to microcircuit function. Science 345, 1255263.

Huang, Z.J., Di Cristo, G., and Ango, F. (2007). Development of GABA innervation in the cerebral and cerebellar cortices. Nat. Rev. Neurosci. 8, 673–686.

Huchzermeyer, C., Albus, K., Gabriel, H.-J., Otáhal, J., Taubenberger, N., Heinemann, U., Kovács, R., and Kann, O. (2008). Gamma oscillations and spontaneous network activity in the hippocampus are highly sensitive to decreases in pO2 and concomitant changes in mitochondrial redox state. J. Neurosci. 28, 1153–1162.

Huchzermeyer, C., Berndt, N., Holzhütter, H.-G., and Kann, O. (2013). Oxygen consumption rates during three different neuronal activity states in the hippocampal CA3 network. J. Cereb. Blood Flow Metab. 33, 263–271.

Inan, M., Zhao, M., Manuszak, M., Karakaya, C., Rajadhyaksha, A.M., Pickel, V.M., Schwartz, T.H., Goldstein, P.A., and Manfredi, G. (2016). Energy deficit in parvalbumin neurons leads to circuit dysfunction, impaired sensory gating and social disability. Neurobiol. Dis. 93, 35–46.

Isokawa, M. (1997). Membrane time constant as a tool to assess cell degeneration. Brain Res Brain Res Protoc 1, 114–116.

Janak, P.H., and Tye, K.M. (2015). From circuits to behaviour in the amygdala. Nature 517, 284–292.

Jonas, P., Bischofberger, J., Fricker, D., and Miles, R. (2004). Interneuron Diversity series: Fast in, fast out--temporal and spatial signal processing in hippocampal interneurons. Trends Neurosci. 27, 30–40.

Kann, O. (2011). The energy demand of fast neuronal network oscillations: insights from brain slice preparations. Front. Pharmacol. 2, 90.

Kann, O. (2016). The interneuron energy hypothesis: Implications for brain disease. Neurobiol. Dis. 90, 75–85.

Kann, O., and Kovács, R. (2007). Mitochondria and neuronal activity. Am. J. Physiol. Cell Physiol. 292, C641–57.

Kann, O., Huchzermeyer, C., Kovács, R., Wirtz, S., and Schuelke, M. (2011). Gamma oscillations in the hippocampus require high complex I gene expression and strong functional performance of mitochondria. Brain 134, 345–358.

Kann, O., Papageorgiou, I.E., and Draguhn, A. (2014). Highly energized inhibitory interneurons are a central element for information processing in cortical networks. J. Cereb. Blood Flow Metab. 34, 1270–1282.

Kann, O., Hollnagel, J.-O., Elzoheiry, S., and Schneider, J. (2016). Energy and Potassium Ion Homeostasis during Gamma Oscillations. Front. Mol. Neurosci. 9, 47.

Kwon, S.-K., Sando, R., Lewis, T.L., Hirabayashi, Y., Maximov, A., and Polleux, F. (2016). LKB1 Regulates Mitochondria-Dependent Presynaptic Calcium Clearance and Neurotransmitter Release Properties at Excitatory Synapses along Cortical Axons. PLoS Biol. 14, e1002516.

Lin-Hendel, E.G., McManus, M.J., Wallace, D.C., Anderson, S.A., and Golden, J.A. (2016). Differential Mitochondrial Requirements for Radially and Non-radially Migrating Cortical Neurons: Implications for Mitochondrial Disorders. Cell Rep. 15, 229–237.

Liu, Q.A., and Shio, H. (2008). Mitochondrial morphogenesis, dendrite development, and synapse formation in cerebellum require both Bcl-w and the glutamate receptor delta2. PLoS Genet. 4, e1000097.

Longair, M.H., Baker, D.A., and Armstrong, J.D. (2011). Simple Neurite Tracer: open source software for reconstruction, visualization and analysis of neuronal processes. Bioinformatics 27, 2453–2454.

López-Doménech, G., Higgs, N.F., Vaccaro, V., Roš, H., Arancibia-Cárcamo, I.L., MacAskill, A.F., and Kittler, J.T. (2016). Loss of Dendritic Complexity Precedes Neurodegeneration in a Mouse Model with Disrupted Mitochondrial Distribution in Mature Dendrites. Cell Rep. 17, 317–327.

López-Doménech, G., Covill-Cooke, C., Ivankovic, D., Halff, E.F., Sheehan, D.F., Norkett, R., Birsa, N., and Kittler, J.T. (2018). Miro proteins coordinate microtubule- and actin-dependent mitochondrial transport and distribution. EMBO J. 37, 321–336.

MacAskill, A.F., and Kittler, J.T. (2010). Control of mitochondrial transport and localization in neurons. Trends Cell Biol. 20, 102–112.

Macaskill, A.F., Rinholm, J.E., Twelvetrees, A.E., Arancibia-Carcamo, I.L., Muir, J., Fransson, A., Aspenstrom, P., Attwell, D., and Kittler, J.T. (2009). Miro1 is a calcium sensor for glutamate receptor-dependent localization of mitochondria at synapses. Neuron 61, 541–555.

Madisen, L., Mao, T., Koch, H., Zhuo, J., Berenyi, A., Fujisawa, S., Hsu, Y.-W.A., Garcia, A.J., Gu, X., Zanella, S., et al. (2012). A toolbox of Cre-dependent optogenetic transgenic mice for light-induced activation and silencing. Nat. Neurosci. 15, 793–802.

Mann, E.O., Suckling, J.M., Hajos, N., Greenfield, S.A., and Paulsen, O. (2005). Perisomatic feedback inhibition underlies cholinergically induced fast network oscillations in the rat hippocampus in vitro. Neuron 45, 105–117.

Marín, O. (2012). Interneuron dysfunction in psychiatric disorders. Nat. Rev. Neurosci. 13, 107–120.

Meijering, E., Dzyubachyk, O., and Smal, I. (2012). Methods for cell and particle tracking. Meth. Enzymol. 504, 183–200.

Montgomery, S.M., and Buzsáki, G. (2007). Gamma oscillations dynamically couple hippocampal CA3 and CA1 regions during memory task performance. Proc. Natl. Acad. Sci. USA 104, 14495–14500.

Morris, R.L., and Hollenbeck, P.J. (1995). Axonal transport of mitochondria along microtubules and F-actin in living vertebrate neurons. J. Cell Biol. 131, 1315–1326.

Motulsky, H.J., and Brown, R.E. (2006). Detecting outliers when fitting data with nonlinear regression - a new method based on robust nonlinear regression and the false discovery rate. BMC Bioinformatics 7, 123.

Myatt, D.R., Hadlington, T., Ascoli, G.A., and Nasuto, S.J. (2012). Neuromantic - from semi-manual to semi-automatic reconstruction of neuron morphology. Front Neuroinformatics 6, 4.

Nguyen, T.T., Oh, S.S., Weaver, D., Lewandowska, A., Maxfield, D., Schuler, M.-H., Smith, N.K., Macfarlane, J., Saunders, G., Palmer, C.A., et al. (2014). Loss of Miro1-directed mitochondrial movement results in a novel murine model for neuron disease. Proc. Natl. Acad. Sci. USA 111, E3631–40.

Nie, F., and Wong-Riley, M.T. (1995). Double labeling of GABA and cytochrome oxidase in the macaque visual cortex: quantitative EM analysis. J. Comp. Neurol. 356, 115–131.

Okaty, B.W., Miller, M.N., Sugino, K., Hempel, C.M., and Nelson, S.B. (2009). Transcriptional and electrophysiological maturation of neocortical fast-spiking GABAergic interneurons. J. Neurosci. 29, 7040–7052.

Pathak, D., Shields, L.Y., Mendelsohn, B.A., Haddad, D., Lin, W., Gerencser, A.A., Kim, H., Brand, M.D., Edwards, R.H., and Nakamura, K. (2015). The role of mitochondrially derived ATP in synaptic vesicle recycling. J. Biol. Chem. 290, 22325–22336.

Paul, A., Crow, M., Raudales, R., He, M., Gillis, J., and Huang, Z.J. (2017). Transcriptional architecture of synaptic communication delineates gabaergic neuron identity. Cell 171, 522–539.e20.

Pelkey, K.A., Chittajallu, R., Craig, M.T., Tricoire, L., Wester, J.C., and McBain, C.J. (2017). Hippocampal gabaergic inhibitory interneurons. Physiol. Rev. 97, 1619–1747.

Pham, A.H., McCaffery, J.M., and Chan, D.C. (2012). Mouse lines with photo-activatable mitochondria to study mitochondrial dynamics. Genesis 50, 833–843.

del Río, J.A., de Lecea, L., Ferrer, I., and Soriano, E. (1994). The development of parvalbumin-immunoreactivity in the neocortex of the mouse. Brain Res. Dev. Brain Res. 81, 247–259.

Rodriguez, A., Zhang, H., Klaminder, J., Brodin, T., Andersson, P.L., and Andersson, M. (2017). ToxTrac : A fast and robust software for tracking organisms. Methods in ecology and evolution / British Ecological Society. doi: 10.1111/2041-210X.12874.

Russo, G.J., Louie, K., Wellington, A., Macleod, G.T., Hu, F., Panchumarthi, S., and Zinsmaier, K.E. (2009). Drosophila Miro is required for both anterograde and retrograde axonal mitochondrial transport. J. Neurosci. 29, 5443–5455.

Sainath, R., Ketschek, A., Grandi, L., and Gallo, G. (2017). CSPGs inhibit axon branching by impairing mitochondria-dependent regulation of actin dynamics and axonal translation. Dev. Neurobiol. 77, 454–473.

Saotome, M., Safiulina, D., Szabadkai, G., Das, S., Fransson, A., Aspenstrom, P., Rizzuto, R., and Hajnóczky, G. (2008). Bidirectional Ca2+-dependent control of mitochondrial dynamics by the Miro GTPase. Proc. Natl. Acad. Sci. USA 105, 20728–20733.

Seibenhener, M.L., and Wooten, M.C. (2015). Use of the Open Field Maze to measure locomotor and anxiety-like behavior in mice. J. Vis. Exp. e52434.

Shlevkov, E., Basu, H., Bray, M.-A., Sun, Z., Wei, W., Apaydin, K., Karhohs, K., Chen, P.-F., Smith, J.L.M., Wiskow, O., et al. (2019). A High-Content Screen Identifies TPP1 and Aurora B as Regulators of Axonal Mitochondrial Transport. Cell Rep. 28, 3224–3237.e5.

Smith, G.M., and Gallo, G. (2018). The role of mitochondria in axon development and regeneration. Dev. Neurobiol. 78, 221–237.

Smith, H.L., Bourne, J.N., Cao, G., Chirillo, M.A., Ostroff, L.E., Watson, D.J., and Harris, K.M. (2016). Mitochondrial support of persistent presynaptic vesicle mobilization with age-dependent synaptic growth after LTP. Elife 5. eLife, 5. doi: 10.7554/eLife.15275.

Sohal, V.S. (2016). How close are we to understanding what (if anything) γ oscillations do in cortical circuits? J. Neurosci. 36, 10489–10495.

Sohal, V.S., Zhang, F., Yizhar, O., and Deisseroth, K. (2009). Parvalbumin neurons and gamma rhythms enhance cortical circuit performance. Nature 459, 698–702.

Spillane, M., Ketschek, A., Merianda, T.T., Twiss, J.L., and Gallo, G. (2013). Mitochondria coordinate sites of axon branching through localized intra-axonal protein synthesis. Cell Rep. 5, 1564–1575.

Stephen, T.-L., Higgs, N.F., Sheehan, D.F., Al Awabdh, S., López-Doménech, G., Arancibia-Carcamo, I.L., and Kittler, J.T. (2015). Miro1 Regulates Activity-Driven Positioning of Mitochondria within Astrocytic Processes Apposed to Synapses to Regulate Intracellular Calcium Signaling. J. Neurosci. 35, 15996–16011.

Stoppini, L., Buchs, P.A., and Muller, D. (1991). A simple method for organotypic cultures of nervous tissue. J. Neurosci. Methods 37, 173–182.

Sun, T., Qiao, H., Pan, P.-Y., Chen, Y., and Sheng, Z.-H. (2013). Motile axonal mitochondria contribute to the variability of presynaptic strength. Cell Rep. 4, 413–419.

Swietek, B., Gupta, A., Proddutur, A., and Santhakumar, V. (2016). Immunostaining of Biocytin-filled and Processed Sections for Neurochemical Markers. J. Vis. Exp. doi: 10.3791/54880

Vaccaro, V., Devine, M.J., Higgs, N.F., and Kittler, J.T. (2017). Miro1-dependent mitochondrial positioning drives the rescaling of presynaptic Ca2+ signals during homeostatic plasticity. EMBO Rep. 18, 231–240.

Wang, X., and Schwarz, T.L. (2009). The mechanism of Ca2+ -dependent regulation of kinesin-mediated mitochondrial motility. Cell 136, 163–174.

Whittaker, R.G., Turnbull, D.M., Whittington, M.A., and Cunningham, M.O. (2011). Impaired mitochondrial function abolishes gamma oscillations in the hippocampus through an effect on fast-spiking interneurons. Brain 134, e180

Zou, D., Chen, L., Deng, D., Jiang, D., Dong, F., McSweeney, C., Zhou, Y., Liu, L., Chen, G., Wu, Y., et al. (2016). DREADD in parvalbumin interneurons of the dentate gyrus modulates anxiety, social interaction and memory extinction. Curr. Mol. Med. 16, 91–102.

